# Disturbed trophoblast transition links preeclampsia progression from placenta to the maternal syndrome

**DOI:** 10.1101/2022.10.10.511539

**Authors:** Olivia Nonn, Olivia Debnath, Daniela S. Valdes, Katja Sallinger, Ali Kerim Secener, Sandra Haider, Cornelius Fischer, Sebastian Tiesmeyer, Jose Nimo, Thomas Kuenzer, Theresa Maxian, Martin Knöfler, Philipp Karau, Hendrik Bartolomaeus, Thomas Kroneis, Alina Frolova, Lena Neuper, Nadine Haase, Kristin Kräker, Sarah Kedziora, Désirée Forstner, Stefan Verlohren, Christina Stern, Fabian Coscia, Meryam Sugulle, Stuart Jones, Basky Tilaganathan, Roland Eils, Berthold Huppertz, Amin El-Heliebi, Anne Cathrine Staff, Dominik N. Müller, Ralf Dechend, Martin Gauster, Naveed Ishaque, Florian Herse

## Abstract

Pre-eclampsia (PE) is a syndrome that affects multiple organ systems and is the most severe hypertensive disorder in pregnancy. It frequently leads to preterm delivery, maternal and fetal morbidity and mortality and life-long complications^1^. We currently lack efficient screening tools^2, 3^ and early therapies^4, 5^ to address PE. To investigate the early stages of early onset PE, and identify candidate markers and pathways, we performed spatio-temporal multi-omics profiling of human PE placentae and healthy controls and validated targets in early gestation in a longitudinal clinical cohort. We used a single-nuclei RNA-seq approach combined with spatial proteo- and transcriptomics and mechanistic *in vitro* signalling analyses to bridge the gap from late pregnancy disease to early pregnancy pathomechanisms. We discovered a key disruption in villous trophoblast differentiation, which is driven by the increase of transcriptional coactivator p300, that ultimately ends with a senescence-associated secretory phenotype (SASP) of trophoblasts. We found a significant increase in the senescence marker activin A in preeclamptic maternal serum in early gestation, before the development of clinical symptoms, indicating a translation of the placental syndrome to the maternal side. Our work describes a new disease progression, starting with a disturbed transition in villous trophoblast differentiation. Our study identifies potential pathophysiology-relevant biomarkers for the early diagnosis of the disease as well as possible targets for interventions, which would be crucial steps toward protecting the mother and child from gestational mortality and morbidity and an increased risk of cardiovascular disease later in life.

## Main

Recent single-cell sequencing studies of healthy female reproductive tissues outside and during pregnancy have characterised the early maternal-fetal interface^6, 7^, trophoblast subtypes^8^, and the endometrium before pregnancy^9^. The temporary maternal-fetal interface, which exists for the duration of pregnancy, mediates *in utero* conditions that facilitate successful pregnancies and also shape the future health of the mother and child over the long term^1^. While the single-cell landscape of healthy placentae has been described well, this is not true for the uteroplacental tissue of those suffering from the hypertensive pregnancy disorder pre-eclampsia (PE). A multi-modal characterisation of PE would provide a deeper understanding of this early gestational pathophysiology and improve clinical management^2, 10, 11^.

Hypertensive disorders in pregnancy account for 14% of maternal deaths^11^. Early-onset pre-eclampsia (eoPE) that urges delivery before the 34^th^ week of gestation is even more destructive^4, 12, 13^. Currently, diagnoses are made based on clinical signs which appear when PE has progressed, late in pregnancy, to a point that maternal and fetal morbidity is often already irreversible^2, 14^ Specific, reliable early first trimester screening methods are lacking^2, 15^. The only pharmacological intervention known to reduce risk, low-dose aspirin, is accompanied by challenges such as insufficient effects in up to 60% of high-risk pregnancies^16^. PE is also associated with a reduction of the lifespan of both mother and child due to complications from cardiovascular disease later in life^2, 10, 11^. Highlighting the fundamental role for the placenta, the only therapy to end the maternal PE crisis, is to deliver the placenta^17, 18^.

Here we integrated single nuclei RNA sequencing (snRNA-seq) with spatially resolved proteo- and transcriptomics and multicentric pre-eclampsia cohorts. We identified an early disruption in the villous trophoblast differentiation transition to the secretory trophoblast (syncytiotrophoblast, STB) lineage and premature senescence that leads to a fetal to maternal syndrome translation in eoPE. The markers and pathways we identifed in the fetal-maternal barrier can be considered as candidates for underlying molecular placental pathology in PE. In turn, these may act as prognostic biomarkers to identify eoPE before clinical symptoms arise, potentially offering crucial tools for the early diagnosis of this serious syndrome as well as therapeutic approaches.

## Results

### Maternal-fetal crosstalk in early and late pregnancy

Since the placenta (fetal tissue) and decidua (maternal tissue) biopsy sampling in ongoing pregnancy increases the risk for adverse outcomes, there is a lack of comprehensive longitudinal studies and understanding of early pathomechanisms translating disease to late gestation. To better understand the early pathophysiology of maternal-fetal crosstalk in eoPE, we analysed diseased preeclamptic and healthy term tissue. In addition, we included early gestation tissue to set pathological findings in a temporal context. We performed snRNA-seq of fetal chorionic villi and maternal decidua in healthy pregnancies from first trimester (5-10 gestational weeks, n=79,885 villus-derived nuclei (v) and 15,367 decidua-derived nuclei (d), in total 95,252 nuclei; early control = e.ctrl; **Fig. 1a-e, Extended Data Table 1 and Extended Data Fig. 1**) and term (≥38 gestational weeks; n=39,663 nuclei; late control = l.ctrl; **Fig. 1a-e**). Additionally, we profiled villi and decidua from women who had developed eoPE (≤34 gestational weeks; n=35,662 nuclei; **Fig. 1a-e**). Single nuclei were harmonised across samples from healthy early and term late, as well as eoPE pregnancies. Differences of eoPE and l.ctrl were computationally adjusted for gestational age differences, since eoPE is defined as PE with pre-term delivery before the 34^th^ week of gestation compared to normal term delivery in l.ctrl. Furthermore, we studied the spatial heterogeneity of cell types by integrating multi-omics data using snRNA-seq, *Visium* spatial transcriptomic assays, spatially resolved *in situ* sequencing (ISS^19^), and spatial proteomics^20^.

**Fig. 1.**
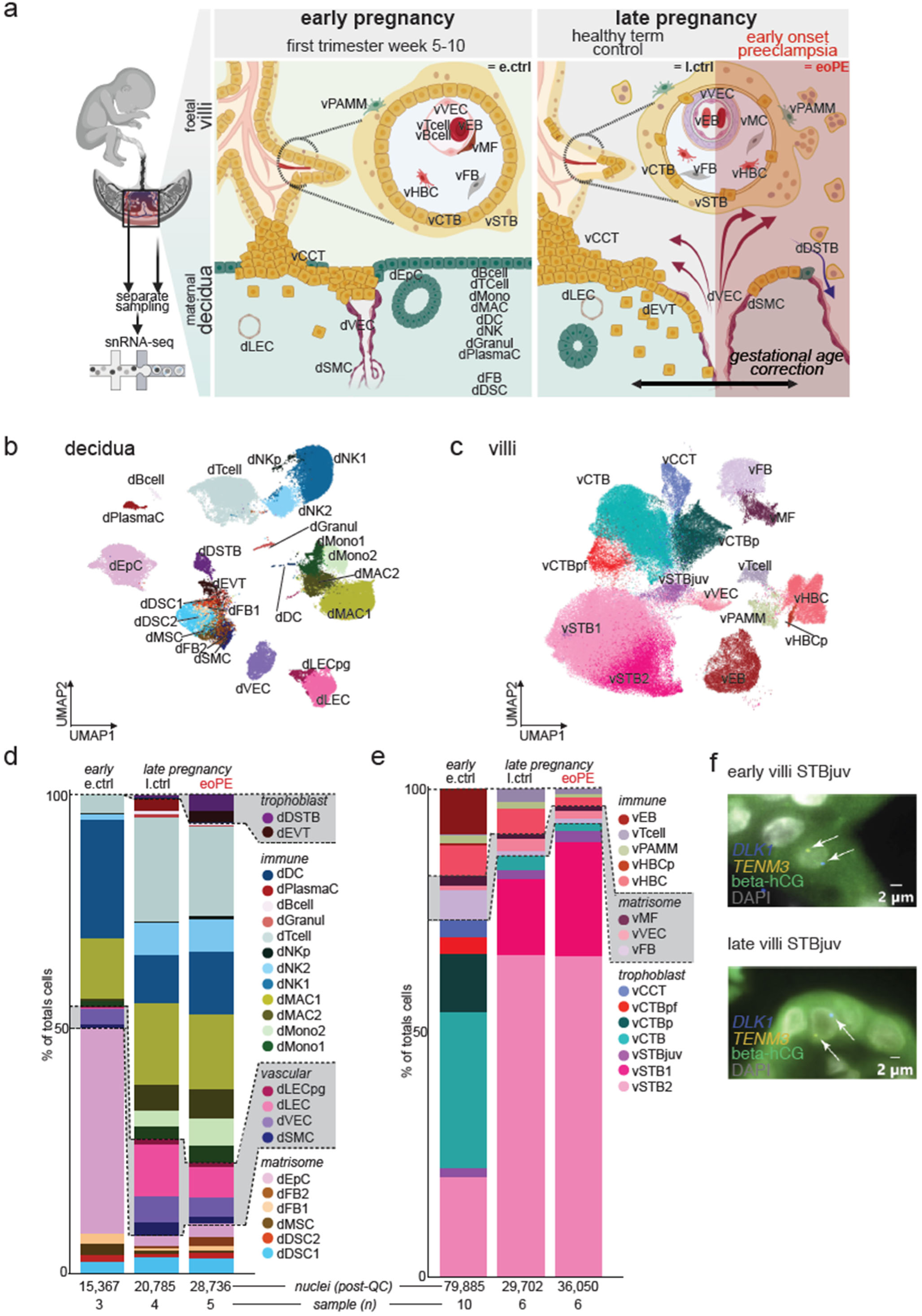
Cell type and state distribution is altered across gestation and disturbed in early-onset pre-eclampsia. **(a)** Schematic illustration of experimental design and histological changes across stages of gestation (early and late pregnancy) and in late pregnancy disease versus control (healthy term=l.ctrl, early onset preeclampsia=eoPE). Placental tissue was separately sampled surgically to collect villi and decidua from the same patients for snRNA-seq. Early tissues correspond to gestational ages between 5-10 weeks (e.ctrl), early-onset pre-eclampsia before 34 weeks (eoPE; range 27-33 weeks of gestation) and late healthy control at 39 weeks (l.ctrl; range is 38-40 weeks of gestation). The gestational age difference between healthy term controls (l.ctrl) and diseased pre-term eoPE were corrected for using additional scRNA-seq data^25^ from preterm controls (pt.ctrl). Cell name abbreviations in Extended Data Fig. 1c. **(b, c)** UMAP displaying **(b)** maternal (decidual, *d*) and **(c)** fetal (villous, *v*) cell types and states from single nuclei RNA sequencing, with integrated samples from e.ctrl, l.ctrl and eoPE. Colours code for a cell type or state. **(d,e)** Cell composition (%) distribution displayed, numbers under bars indicate the sample size of sequenced nuclei. Cell compositions presented across gestational time points (e.ctrl, l.ctrl) and between disease states (eoPE) are for **(d)** decidua and **(e)** villi. **(f)** Representative images showing the localisation of the novel STB cell state STBjuv; immunofluorescence staining with STB protein marker βHCG (green) combined with padlock probe based *in situ* hybridisation for STBjuv markers *DLK1* and *TENM3* (arrows indicate STBjuv specific mRNA markers). Positive and negative controls for the probes are shown above. STB, syncytiotrophoblast; juv, juvenile state. n = 3 independent experiments with 2 biological replicates each per gestational time.

We identified a rich diversity of cell types and cell states in the maternal-fetal barrier (**Fig. 1c, Extended Data Fig. 1**). Variations in cell composition within the immune, vascular-endothelial, matrisome, and trophoblast compartments were evident at different gestational sampling times at the maternal (**Fig. 1d, Extended Data Fig. 1 and Table 4**) and fetal (**Fig. 1e, Extended Data Fig. 1 and Table 3**) interface, mirroring specific functional adaptiations at different stages of pregnancy. Syncytiotrophoblast (STB) populations were more prominent in late compared to early villi, and they tended to be more abundant in eoPE tissues (percent of total nuclei: 22.2% in e.ctrl, 83.3% in l.ctrl and 91.2% in eoPE, FDR < 0.01 e.ctrl to l.ctrl, FDR=0.144 l.ctrl to eoPE). We identified nuclei in the decidua that are villous-derived and strongly positive for STB markers such as *CGB* and *KISS1*. We defined them as “deported STBs” found in decidua (dDSTB). These fragments released from the STB are found more frequently in eoPE decidua than in controls (eoPE= 980; l.ctrl=85, e.ctrl=0; **Fig. 1a, b, e Extended Data Table 4**), and are postulated to be the final product of senescent vSTB when shed into maternal circulation^21, 22^.

Our integrative analysis of snRNA-seq and spatial data showed that STB exists in three transcriptionally different nuclei states, including a novel STB juvenile subtype (vSTBjuv) alongside vSTB1 and vSTB2 (**Fig. 1c, e** and **f, Extended Data Fig. 2**). The vSTBjuv population is characterised by the notably higher expression of hormone genes including placental lactogen (*CSH1, CSH2*). It also exhibits exocytotic expression signatures (*HSPB1, CD63, FURIN)* in addition to more classical vSTB genes: *KISS1, CGA, PGF, EBI3, TFPI*. We postulate that vSTBjuv has a stromal function by regulating cytoskeletal stability and the extracellular matrix, as indicated by the expression of genes such as *ACTB, TMSB10, SPARC,* and *VIM* (**Extended Data Fig. 2**).

Next, we localised the vSTBjuv cell state within the STB layer. We identified markers that best distinguished vSTBjuv from vSTB1 and vSTB2, which were *TENM3* (log2FC=12.4), which promotes homophilic adhesion^23^, and *DLK1* (log2FC=6.2), a paternally imprinted gene which is correlated with birthweight^24^. Using *in situ* hybridisation with *DLK1* and *TENM3* padlock-probes, we localised *DLK1*^+^/*TENM3*^+^ vSTBjuv within a β-hCG^pos^ STB layer (**Fig. 1f**; **Extended Data Fig. 3**). We validated the transcriptomic signatures of the STB and their progenitors, the cytotrophoblasts (CTBs), which are both components of the maternal-fetal barrier, by applying spatial proteomics^20^ (**Fig. 2b-c**, **Extended Data Fig. 4a-d**). The revealed signatures showed enrichment for signalling of RhoGTPases, MIRO and RHOBTB3, and enriched mitotic pathways in vCTBs. vSTBs exhibited an enrichment in oxidative stress-associated processes such as ROS production and enzymes known to be involved in metabolic disorders of biological oxidation (**Extended Data Fig. 4**).

**Fig. 2.**
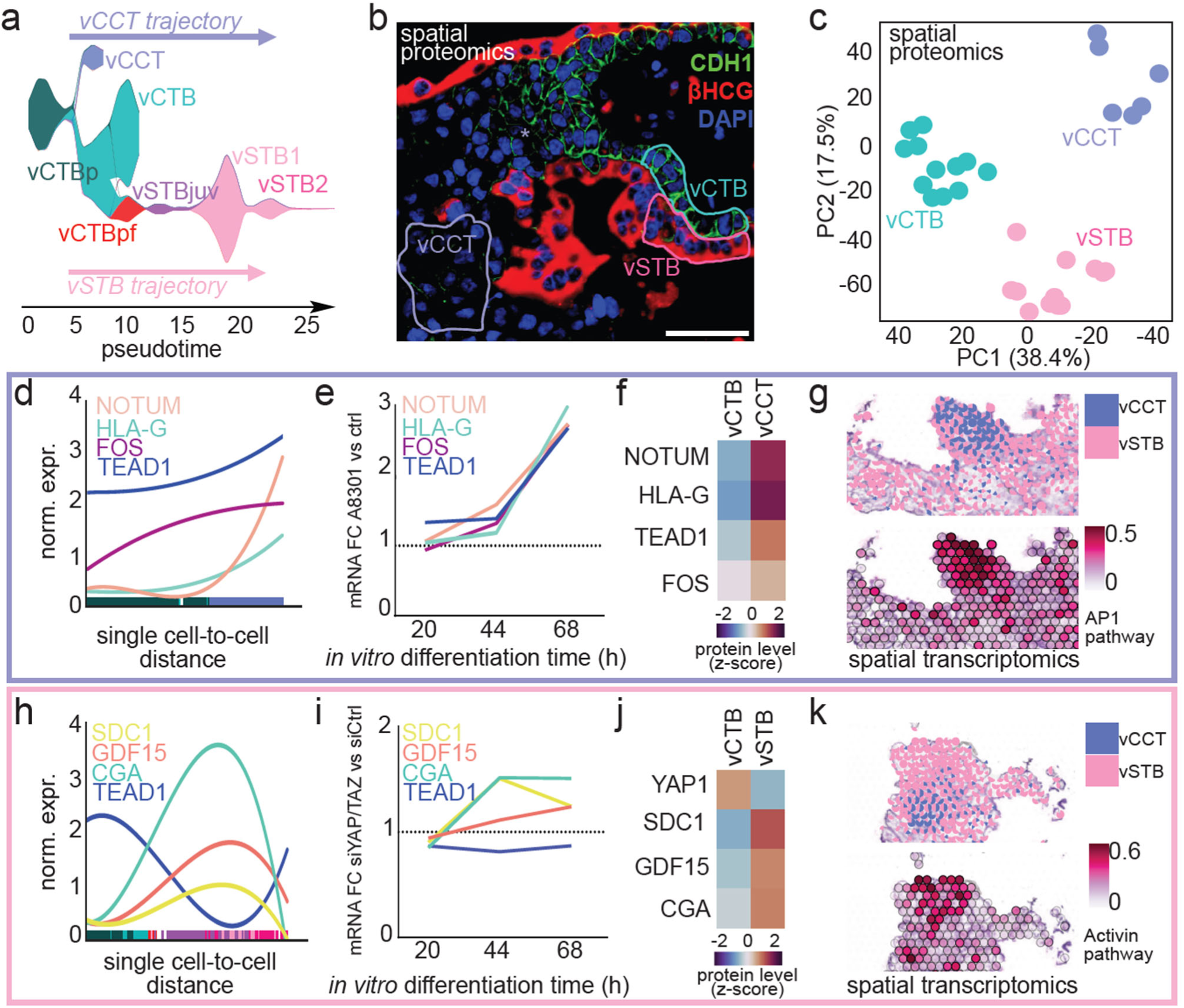
Early trophoblast invasion is AP1 and Hippo pathway driven and early syncytiotrophoblast differentiation is prematurely initiated with silenced YAP/TAZ. **(a)** Stream plot elucidating the developmental trajectory of early trophoblast and cell density across pseudotime. Branch length represents pseudotime progression, branch width is directly proportional to cell numbers at a given pseudotime. **(b)** Early placenta FFPE sections (n=4, gestational age 7-10 weeks) stained using immunofluorescence markers CDH1 and β-hCG; CTB, STB, and CCT were identified and set areas laser micro-dissected. Dissected areas were captured and processed via LC-MS. **(c)** Principal component analysis of proteomics data revealed cell-type resolved proteomes based on 4,403 quantified protein groups. PC1 and PC2 represent 38.4% and 17.5% of the total variability, respectively. **(d)** Trophoblast lineage commitment regulators with a dynamic expression that correlates with pseudotime. Lines are the polynomial regression fits to the normalised gene expression data. Cell-type membership is incorporated on the x-axis, colours correspond to those annotated in Fig. 1b and 2a (snRNA-seq, n=10 early placentae). **(e)** Primary CTB isolated from first trimester placenta were cultured as indicated on x-axis (time in hours). Dynamic genes in vCCT lineage as calculated in snRNA-seq analyses are replicated in CTB incubated with TGFβ-inhibitor A8301 for up to 68 hours (5 µM, n=5 placentae pooled with n=4 independent experiments). **(f)** Spatial proteomics results showing relative protein levels (z-score) of transcriptionally dynamic genes; transition markers NOTUM, HLA-G are highly expressed in CCT, transition gene TEAD1 increases in CCT. Member of AP1-pathway *FOS* expression is strongly increased in vCCT. **(g)** Spatial transcriptomics showing AP1-pathway genes *FOS, JUN, FOSL1* expressed in a cell column as found in non-negative matrix factorisation (NMF) deconvolution based *Visium* analysis, integrating snRNA-seq data. **(h)** Trophoblast lineage commitment regulators with a dynamic expression that correlates with pseudotime. Lines are the polynomial regression fits to the normalised gene expression data. Cell-type membership is incorporated on the x-axis, colours correspond to those annotated in Fig. 1b and 2a (snRNA-seq, n=10 early placentae). **(i)** Primary CTB isolated from first trimester placenta were cultured as indicated on x-axis (time in hours). Dynamic genes in STB lineage as calculated in snRNA-seq analyses are replicated in siYAP/TAZ treated CTB incubated for up to 68 hours (n=5 placentae pooled with n=4 independent experiments). **(j)** Spatial proteomics validate dynamic genes found on a sn-transcriptomic level; STB genes *CGA, SDC1, GDF15* are highly expressed in STB, transition gene YAP decreases in STB. **(k)** Spatial transcriptomics showing Activin-pathway associated genes expressed in deconvoluted STB areas (spots) as found in NMF deconvolution based *Visium* analysis integrating snRNA-seq data. Cell name abbreviations in **Extended Data Fig. 1c**.

### Activin and Hippo reciprocally drive physiological trophoblast transition

To resolve the trophoblast cell states and infer how their disturbed development may play a crucial role in eoPE, we recapitulated the transcriptional dynamics of human trophoblast differentiation from the progenitors CTBs to their final identities cell column trophoblasts (CCTs) or STBs. Modelling cell trajectories using pseudotime with proliferating vCTB (vCTBp) as the root, we detected two distinct lineages of this bipotent trophoblast. The trajectory expressed gene patterns computationally that predicted to be *transition genes* expressed along a trajectory branch. vCTB cell fate ran towards invading vCCT, or towards a secretory vSTB lineages **(Fig. 2a, Supplementary Table 3).**

In the CCT trajectory (**Fig. 2a**), vCTBs can commit towards vCCT^26, 27^, a cell type that expands and proliferates in the proximal part of cell columns (**Fig. 2b**, asterisk). Distally, it anchors villi on maternal decidua. These distal CCT^26, 27^ (**Fig. 2b**, area circled in blue) then migrate and invade maternal decidua to remodel maternal vessels and partially replace maternal local endothelium (called decidual extravillous trophoblasts = dEVTs, **Fig. 1b**). This so-called spiral artery remodelling results in low-flow, low-resistance vessels that prevents damage to the fetal trophoblast barrier. A faulty spiral artery remodelling initiated by vCCT has been postulated as a crucial event in the development of PE^14, 28^. Receptor-ligand analyses of vCCT prior to invasion and non-invaded maternal decidua suggested that vCCT may initiate maternal decidua reorganisation via several integrins (*ITGB8*, *ITGA1*), *FLT1*, and *TGFBR1* (**Extended Data Fig. 4e**). Key transition genes in the transition of vCTB towards the vCCT phenotype are *HLA*-*G* and *NOTUM* (**Fig. 2d and Extended figure 4f**), so we investigated potential drivers of the vCCT phenotype. We observed dynamic increase of transcription factors *FOS* and the Hippo pathway member *TEAD1* along the pseudotime axis (**Fig. 2d**), which reflected the change from epithelial-like CDH1^pos^ vCTB to vCCT, that are ready to migrate and invade. The YAP/TAZ-TEAD1 complex, which are Hippo pathway members, can act with other transcription factors with distinct functional outputs^27^. *TEAD1* in combination with *FOS* is known to trigger epithelial to mesenchymal transition and migration^29, 30^, a prerequisite for vCCT driven maternal tissue remodelling. Inhibition of activin/nodal receptors ALK4/5/7 is known to shift the vCTB progenitors into vCCT and dEVT transition^31^. We validated the transition to *HLA*- *G*^+^ vCCT in primary isolated first trimester CTB by inhibiting ALK4/5/7 with A8301 and thus recapitulated that vCCT transition genes *HLA-G, NOTUM, TEAD1* and *FOS* increased over time (**Fig. 2e**). To confirm these findings at the protein level, we identified vCTB, vSTB, and vCCT on FFPE sections, laser microdissected the cell type specific areas and then performed LC-MS (**Fig. 2b**). Principal component analysis and unsupervised hierarchical clustering showed that vCTB was observed between terminal differentiation states vCCT and vSTB on the proteome level (**Fig. 2b-c, Extended Data Fig. 4b, d**). Our quantitative protein analyses revealed a significantly higher expression of FOS in combination with higher TEAD1 in HLA-G^+^/NOTUM^+^ vCCT (**Fig. 2f**). Using *Visium* spatial transcriptomics to replicate our previous findings, we saw high expression of the AP1 pathway genes *FOS*, *FOSL1*, and *JUN* specifically localised in vCCT regions (**Fig. 2g**), further highlighting that AP1 signalling contributes to the development of vCCT with their migratory and invasive phenotype^32, 33^.

Next to the CCT trajectory, we also investigated differentiation dynamics of the vSTB lineages (**Fig. 2h-k**). It has been postulated that the fetal placental villi, namely the secretory multinucleated vSTB cell type covering villi as a barrier to maternal blood, secretes factors into maternal circulation causing PE as maternal syndrome^34, 35^. In the vSTB trajectory, *ERVFRD1*^+^ pre-fusion CTB (vCTBpf) were defined as representing a dynamic state of transition between vCTB and the novel vSTB cell state vSTBjuv (**Fig. 2a**). vCTB lineage leaf genes such as Hippo pathway effector transcriptional co-activators *YAP1* **(Extended Data Fig. 5**) in combination with *TEAD1,* are important in maintaining proliferation in vCTB^36–38^. During the vSTB transition, *TEAD1* is repressed contrary to *TEAD1* upregulation in vCCT trajectory (**Fig. 2h**).

In agreement with transition genes and pathways identified through our snRNA-seq pseudotime analysis (**Extended Data Fig. 5**), the *in vitro* silencing of Hippo effectors YAP/TAZ (siYAP/TAZ) in primary CTBs prevented *TEAD1* increase (**Fig. 2i**). Our proteomics analysis confirmed that vSTBs had lower levels of YAP protein than vCTB (**Fig. 2j**). Blocking activin receptors ALK4/5/7 is known to contribute to self-renewal of vCTB^29^ similarly to Hippo members YAP^37^. Instead, vSTB express the key transition genes *SDC1*, *CGA*, and the activin-pathway ligand *GDF15*^39^ (**Fig. 2h**). Consistent with transition genes, silencing YAP/TAZ in primary vCTB provoked an increase in the expression of *SDC1*, *CGA*, and *GDF15* over time (**Fig. 2i**). Spatial transcriptomics, on the other hand, showed an increase of TGFβ-related placental BMP pathway related genes such as *BMP1, SMAD6, GREM2* in vSTB-specific deconvoluted regions (**Fig. 2k**).

In summary, Hippo^37^ and activin^31^ pathways are crucial for maintaining CTBs and an adequate differentiation towards AP1 pathway enriched vCCT^26^ as well as vSTB^37^. Both trophoblast cell types form the interface and barrier between fetal and maternal circulations. Our multi-omics *in situ* and *ex vivo* data replicates previous *in vitro* findings^37^ that a loss of Hippo drives CTB-STB transition towards activin-enriched nuclei states of secretory vSTBs, while activin inhibition drives migratory CCT differentiation by combining the Hippo and AP1 pathways.

### vSTB fusion trajectory is dysregulated in PE

Having established physiological trophoblast transition on a spatial multi-omics level, we next investigated the impact of the pre-eclamptic disease state on the placental transcriptome and the transition processes. Comparisons of eoPE to l.ctrl samples were adjusted for the effect of preterm birth, i.e. the gestational age difference, using published single-cell data specifically characterising differences on non-eoPE pre-term placental cells^19^. For the analysis pipeline, we also adjusted for potential confounders and importantly, we validated the snRNA-seq targets in a multicentre pre-eclampsia cohort.

Since uteroplacental tissue analyses of preeclampsia are limited to late timepoints in pregnancy, we aimed to infer late gestation pathological profiles by computationally recapitulating early gestation pathophysiology and evaluating disturbance. Therefore, we investigated the fusion dynamics of STB progenitor cells with the goal to identify at which stage of eoPE cells depart from their normal developmental trajectory. These STB progenitor cells are *ERVFRD-1*^+^ CTB cells, previously described by Liu et al.^8^ (**Fig. 3a**, here: CTBpf). Notably, CTBpf was the only villous cell type that expressed the BMP-inhibitor *GREM2* (**Extended Data Fig. 2, 8**), and we showed that GREM2^pos^ cells still expressed the classical CTB-marker CDH1 in early villous tissue (**Fig. 3b, c**). The CDH1^pos^ cell borders of CTBpf also show they are not yet fused with the multinucleated vSTB and are still a mono-nucleated vCTB subtype, which is why we named them CTBpf for their pre-fusion CTB cell state (**Fig. 3d**).

**Fig. 3.**
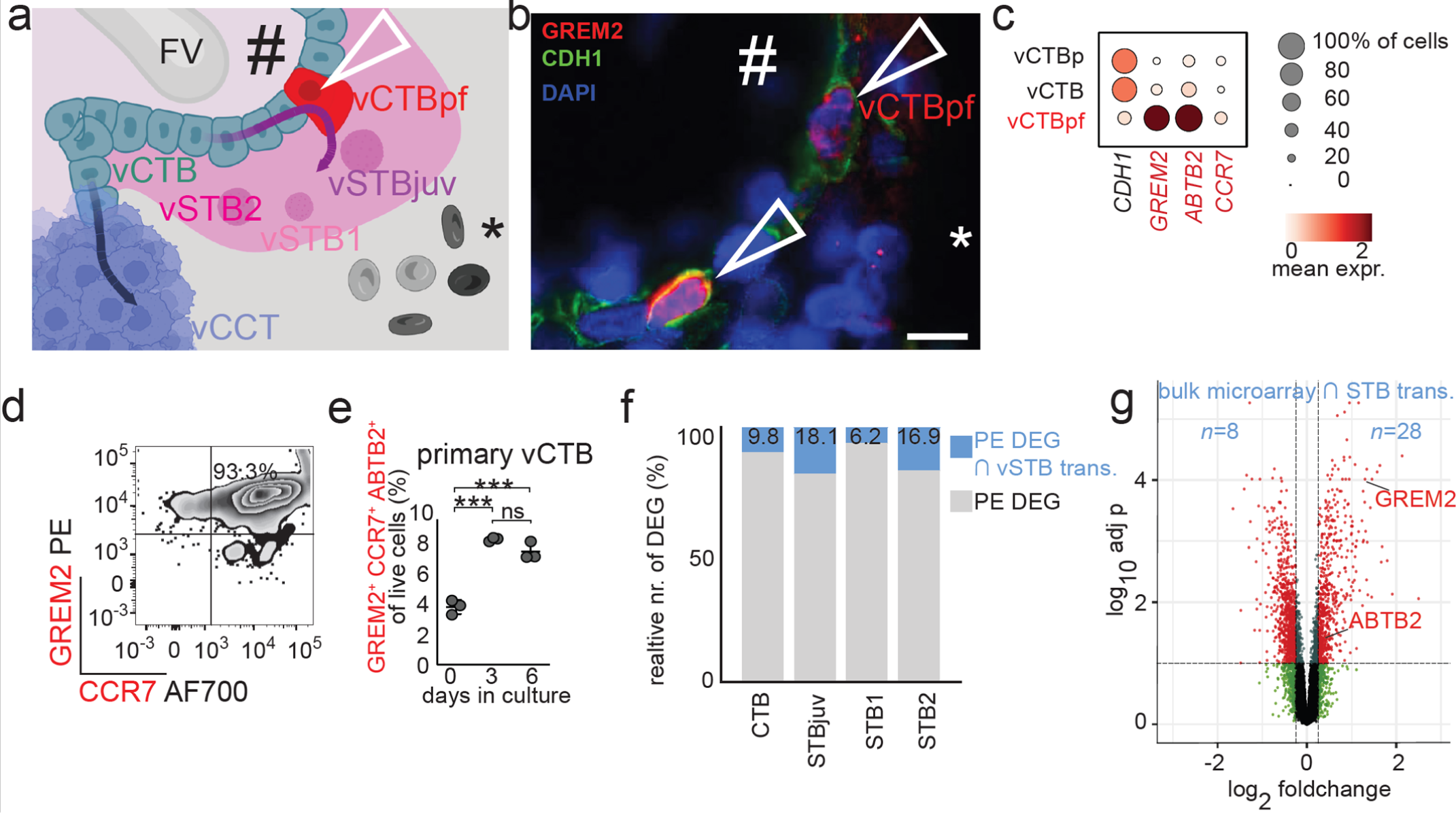
Syncytiotrophoblast pre-fusion cytotrophoblast are dysregulated in early-onset pre-eclampsia. **(a)** Schematic drawing illustrating cell trajectories as found in Fig. 2a; arrow pointing to CTBpf, pre-fusion CTB; FV = fetal vessel, # = villous stroma; * = maternal blood in intervillous space; Cell name abbreviations in Extended Data Fig. 1b. **(b)** Immunofluorescence staining of GREM2^pos^ CTBpf (arrow) within a CDH1^pos^ (E-Cadherin) CTB layer. # = villous stroma; * = maternal blood in intervillous space. **(c)** Dotplot of snRNAseq data with markers used in flow cytometry for CTBpf characterisation. **(d)** Flow cytometry of primary isolated first trimester CTB showing gating strategy; >93% CD49f-/CDH1+ CTB were positive for CTBpf markers GREM2, CCR7, ABTB2. **(e)** Primary first trimester CTB were cultured for 3 and 6 days; a stable increase to ∼8% of GREM2+ CCR7+ ABTB2+ fraction of live cells can be seen (n=3, villous CTB isolated from n=5 placentae and pooled). **(f)** Relative number of differentially expressed genes (DEGs) in CTB and STB states in eoPE. Overlap with transition genes derived from the trophoblast trajectory analysis towards the STB lineage is highlighted in blue, with the exact percentage contribution written on the stacked bar plot. **(g)** Volcano plot of villi dysregulated genes in early-onset PE analysed from a published microarray dataset^40^, highlighting concurrence with key genes identified in this manuscript; each dot represents an individual gene. Interestingly, we observe *GREM2* (vCTBpf state marker and a BMP antagonist) as one of these candidate genes, supporting the notion that dysregulated STB differentiation in eoPE is a mechanistic driver of placental dysfunction. (n = 23 placenta control, 14 eoPE). pt.cntrl; preterm control, eoPE; lateC, late pregnancy control; eoPE, early onset pre-eclampsia. STB, syncytiotrophoblast; CTB, cytotrophoblast; pf, pre-fusion.

CTBpf also show high expression of *CTNNB1* mRNA that encodes β-catenin. Spatial proteomics of the STB layer confirmed the presence of β-catenin (**Supplemental Table 9**), a Wnt regulator that is involved in CDH1 degradation. The appearance of *CTNNB1* in CDH1^pos^ fusion-competent CTBpf cells underlines the importance of CTBpf in the transition from CDH^pos^ vCTB to CDH^neg^ vSTB. We performed flow cytometry with primary CDH1^pos^ CTB isolated from first trimester placentae to corroborate these findings. We used snRNA-seq to identify the vCTBpf markers *GREM2, ABTB2, CCR7* (**Fig. 3c**) that we then used in flow cytometry to identify and validate the vCTBpf. >93% of CDH1^pos^/CD49^neg^ were positive for CTBpf markers GREM2 and CCR7 (**Fig. 3d, e; Extended Data Fig. 6f**). This CTBpf fraction increased over time in culture, achieving a stable fraction of around 7-9% of live cells after day 3 (**Fig. 3e**). This was accompanied by a shift from G0/G1 phase to G2/M phase and an increase of DNA content (Extended Data **Fig. 6g)**. These data support our proposal that the “pre-fusion CTB” cell state^8^ is a vCTB subtype and that, as cultured primary CTBs undergo spontaneous fusion, vSTB have passed this vCTBpf cell state. Based on the specific expression of BMP-inhibitor GREM2 in vCTBpf, we validated *in vitro* that BMP7 inhibited fusion induced by cAMP agonist forskolin (**Extended Data Fig. 8**). Temporary BMP inhibition by CTBpf seems to be a prerequisite for fusion from vCTBs to vSTB. This led us to investigate the activity of genes predicted to play key roles in modulating the development of STB and lineage commitment that starts with the CTBpf cell state. We identified a dysregulation of transition genes that drive trophoblast development (**Supplementary Table 3**). STBjuv show the highest percentage of dysregulated STB transition genes (18.1%, **Fig. 3f**) such as *GDF15* and *LNPEP* (**Extended Data Fig. 5**). In comparison to vSTBs, vCTBs have a relatively low fraction of transition genes within their DEG (13 transition genes of 349 total CTB DEG, 0.04%). Notably, *GREM2* was also highly upregulated in the microarray validation dataset of placenta whole tissue (**Fig. 3g**). Herewith, we can indicate that GREM2^pos^ pre-fusion vCTBs are of importance in the aberrant trophoblast differentiation in eoPE. This CTBpf state was also more abundant in first trimester compared to late gestation (**Extended Data Table 3**), and therefore may be part of eoPE pathophysiology already in early pregnancy by impacting vSTB transition.

To make an assessment of the extent of global tissue dysregulation, we measured the number of differentially expressed genes (DEGs) in eoPE compared to term control placentae. We found that between villi and decidua, villous cell types are the most profoundly disturbed (**Fig. 4a, Supplementary Table 4**). Of the global tissue dysregulation, STB accounted for 58.2% of the total DEGs in villi (% of villous DEG, **Supplementary Table 5**). Of note, we found similar patterns in a larger microarray datasets^40^ of eoPE vs control (*n*=37; **Extended Data Fig. 7**), confirming our data in a larger cohort. We found some shared DEG between different villous cell types (**Extended Data Fig. 7**), speculating global dysregulating events. Interestingly, the gene *FLT1* encoding for the anti-angiogenic factor sFLT1 used in the clinic for short-term follow-up in cases of eoPE^3^, was upregulated in two cell types, the vSTB and the vVEC. In summary, eoPE is characterised by a massive dysregulation of the fetal barrier consisting of mainly vSTB, as observed in the high degree of dysregulation in these cell states.

**Fig. 4.**
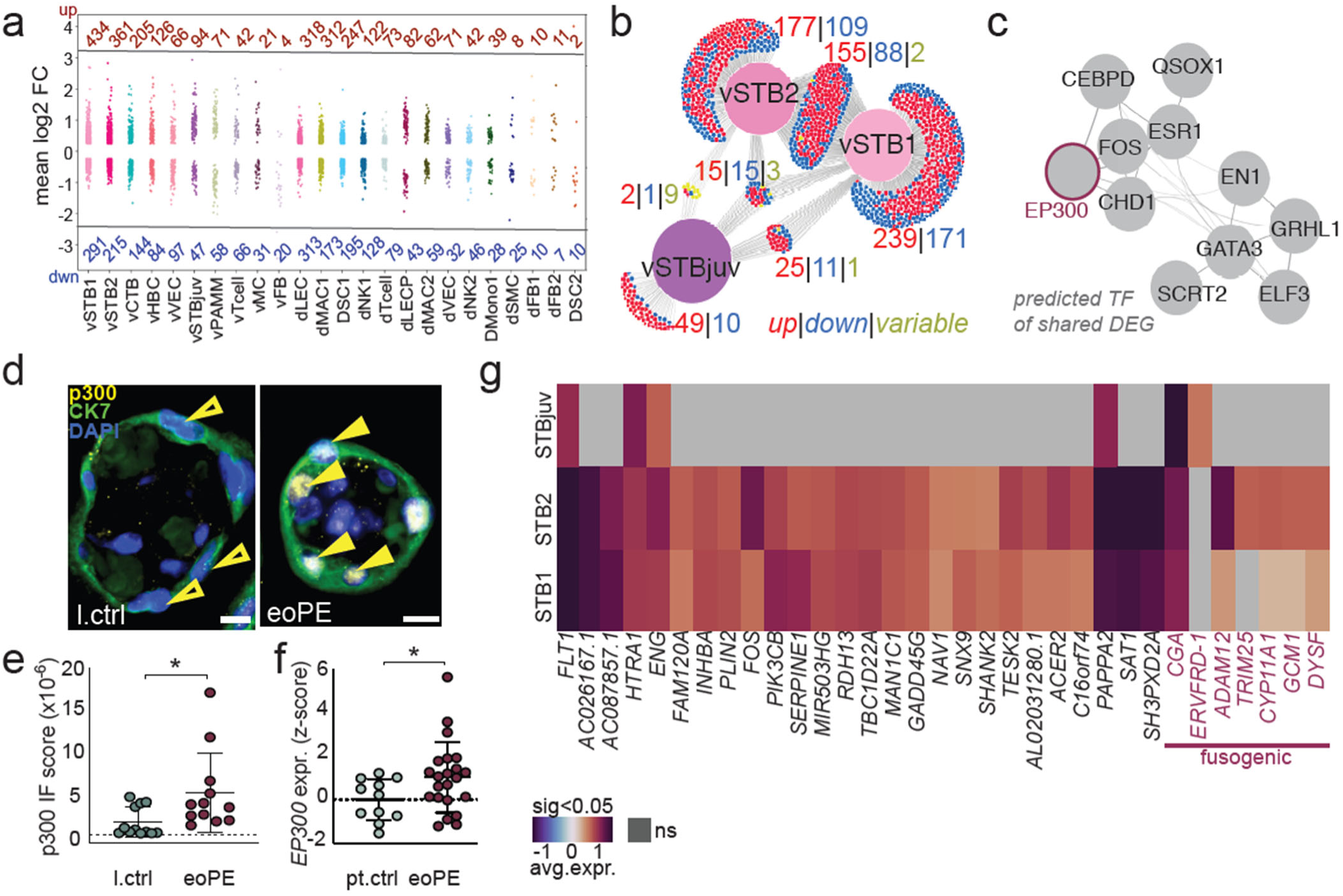
Syncytiotrophoblast populations are most affected in early-onset pre-eclampsia. **a)** Significantly dysregulated expression profiles between late control (l.ctrl) and eoPE villous and decidual cell types (Bonferroni adjusted two-sided, logistic regression p<0.05 and log2FC>± 0.25). Log2FC between conditions for each individually expressed genes (dot) is visualised. vCTBpf, vCCT, vCTBp, vEB, dBcell, dEpC, dNKp, dPC, dGranul, dEVT, dDSTB excluded due to large cell-type composition changes between l.ctrl and eoPE. n=9 deciduas (4 late control, 5 eoPE), n=12 placentae (6 late control, 6 eoPE). The analysis pipeline was adjusted for potential confounders such as the sample collection site, fetal sex, and chemistry used in sequencing. Since the groups did not differ in terms of BMI, maternal age, or smoking habits, we did not make adjustments in these parameters, to avoid overadjustment. **(b)** Convergence of DEGs in the syncytiotrophoblast lineage where each dot represents a single gene (shared upregulated: red; shared downregulated: blue; variable gene expression: green). **(c)** Predicted transcription factors (pTF) were calculated using dysregulation network of trophoblast functional overlap, where motifs of dysregulated shared STB-genes were used to predict its upstream transcription factors (threshold NES >3, max FDR on motif similarity 0.001); *EP300* has most targets within the dysregulated genes (*EP300* marked in red; dysregulated in ≥ 2 STB groups, minimum logFC ≥ 0.4, gene expressed in ≥ 30% cells; n= 28, marked in black). **(d)** p300 is encoded by *EP300*, and was stained in tissue sections of healthy late term controls (l.ctrl, n=3) and eoPE (n=3) where in CK7^pos^ trophoblasts p300 activation through translocation to the nucleus was observed (open arrowheads show nuclei without p300 staining, closed arrowheads show nuclei with p300 staining). **(e)** p300 activation was systematically analysed using automated image analysis by calculating a score normalising positive p300 nuclei to total number of nuclei (calculated via mean intensity values based on immunofluorescence staining of p300 and DAPI, total trophoblast area and overall villi area (calculated via mean intensity values based on CK7 immunofluorescence staining area), dividing each image scanned at same exposure times per channel into quadrants; Wilcoxon rank-sum test, sig.-level 0.05. **(f)** mRNA expression level of *EP300* is significantly increased in eoPE compared to preterm controls (no gestational age difference) in a multicentre cohort, sig.-level 0.05. **(g)** Heatmap illustrating fold changes in log2FC of *EP300* dysregulated targets between eoPE and term controls groups for overlapping genes shown in (b) for STB n=12 villi (6 late control, 6 eoPE), genes involved in fusion are marked in red. Logistic Regression was used for differential testing (log2FC ≥ 0.25 and p-value < 0.05 after Bonferroni adjustment for multiple testing of states, fusogenic *EP300* target genes are marked in red).

vSTB is a multinucleated fused cell barrier towards the maternal circulation and showed the highest aberration (**Fig. 4a**), so next we investigated dysregulation patterns of different STB states and analysed the DEGs shared between STB subtypes (22.6%; n=329 of 1084; **Fig. 4b**). Of these, 99.5% (n=15) showed same directionality in gene expression, indicating a functional unit between vSTB1, vSTB2, and vSTBjuv states. We therefore carried out a regulatory analysis to map transcription factors (TFs) to binding motifs in downstream dysregulated targets shared between vSTB states (**Fig. 4c**) to investigate the drivers of transition of the vSTB lineage under pathological conditions. A TF that emerged from our prediction model as master regulator and top-ranked TF was *EP300*, a transcriptional coactivator associated with fusion^41^ and cell cycle arrest^42^, that has binding motifs in the highest number of STB genes (normalised enrichment score=3.54, FDR < 0.05, n= 561, i.e., 52% targets). In summary, we identified *EP300* as hub TF with the highest number of targets within the DEGs shared between STB states (**Fig. 4c**), and an involvement in fusion processes.

Congruently, we identified an overlap between EP300 target and fusogenic genes including CTBpf marker *ERVFRD-1* and *GCM1,* which are associated with fusion of cells^8^ (**Fig. 4g, Extended Data Fig. 6d**). Dysregulated EP300 target genes in eoPE were also found expressed early in pregnancy and are involved in the STB transition (**Extended Data Fig. 6d, e**)*. EP300* encodes the histone acetyltransferase p300, which is translocated to the nucleus upon activation. We show that the nuclear localisation of p300 in trophoblasts is higher in eoPE compared to l.ctrl (**Fig. 4d, e**). *EP300* was also found to be upregulated in whole-placenta lysates in eoPE compared to gestational age-matched preterm controls (pt.ctrl) in a multicentre cohort (**Fig. 4f**). This suggests that the cause of dysregulation in eoPE may be the aberrant transition to STB, driven by *EP300.* This may subsequently cause an early differentiation or disproportionate shift towards the vSTB-lineage in eoPE.

Altogether, we identified p300/EP300 as important driver in eoPE. Dysregulation in eoPE can be linked to the early transition of STB, at the intermediary fusion states CTBpf and STBjuv.

### eoPE is associated with a senescence-associated secretory phenotype (SASP) at the maternal-fetal interface

Our analyses suggest that eoPE is the result of a perturbation in the transition of STB via fusion cell states. The differentiation trajectory of these cells inherently involves senescence-associated processes^21, 22^. When comparing the DEGs of STB subtypes to the senescence-associated secretory phenotype (SASP) atlas database^43^, we find that in eoPE, affected trophoblasts exhibit a higher increase in SASP gene activity than in late controls. 12% of the genes dysregulated in STBs during eoPE are annotated as SASP factors (**Supplemental Table 7**). The placenta is known to be essential in the pathology of PE and the placental syndrome eventually translates into a maternal systemic syndrome^17^. Altogether, this led us to carry out an analysis of focused receptor-ligand interactions that might affect fetal to maternal SASP translation. Here, we turned to a model that could be used as a proxy for the fetal to maternal crosstalk from trophoblast to the maternal vasculature that underwent changes during pregnancy. To this end, we analysed cell-cell communication between villous derived STB components and decidual vascular cells (dVEC, dSMC) to investigate how the dysregulated STB subtypes we described above could translate the fetal villous syndrome to the maternal system.

We defined anatomically relevant cell types which physiologically interact, and computationally analysed them for their ligand-receptor interaction (**Fig. 5a**). Among the upregulated factors, we found a predominance of secreted factors (of dysregulated vSTB genes were identified as secreted or extracellular exosome factors (GO:0005615, GO:0070062, GO:0032940; GO:0060627; GO:0048010)) that interact with receptors on the maternal vasculature (dVEC, dSMC) via the maternal circulation **Fig. 5a)**. The interactions between vSTB-dVEC and vSTB-dSMC predominantly involved SASP genes (67%, 6 out of 9). *GDF15-TGFBR2* is found to be interacting via maternal blood between villi in vSTB and decidual endothelial cells (**Fig. 5a**). GDF15 is a secreted TGFβ superfamily protein and activin ligand that is abundant in the placenta^44^. In PE, it is observed in increased levels in maternal serum and placentae^45^. In addition, we demonstrated a significant upregulation of vSTB-derived ligands of *LEP-* and *INHBA* as well as interactions with their respective receptors **(Fig. 5a**). Upregulation of both *GDF15* and *INHBA* was independently validated in the multicentre eoPE cohort, where; they were significantly upregulated in patients compared to gestational with age-matched pre-term controls (**Fig. 5b**). Additionally, GDF15 was detected by immunofluorescence to be elevated in eoPE patients compared to healthy controls (**Fig. 5e**). When we visualised all DEGs of vSTBs that were *EP300*-mediated SASP genes, we observed significant aberration profiles of vSTB cell states in eoPE (**Fig. 5f**). We robustly validated our findings using multiple lines of evidence (**Fig. 5b-e**). We conclude that overall, SASP drives the majority of fetal-to-maternal ligand-receptor interactions in eoPE via secretion into the maternal circulation.

**Fig. 5.**
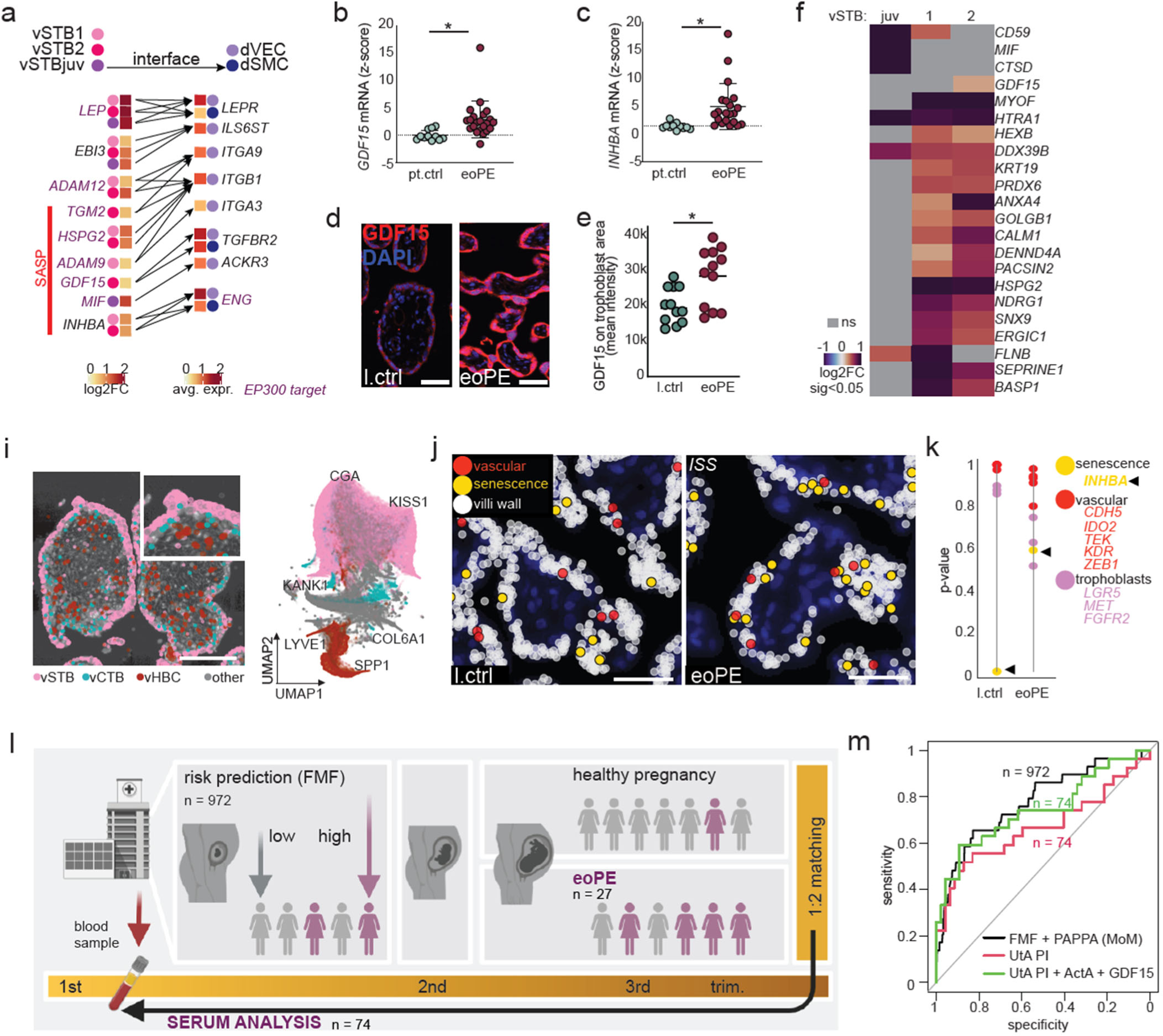
Early-onset pre-eclampsia affects key regulatory interactions at the maternal-fetal interface. Cell crosstalk dysregulation during eoPE in trophoblast secreted ligands. **(a)** vSTB secreted ligands altered in eoPE relative to term controls with preterm gestational age correction acting on highly expressed decidual endothelial receptors highlight ligand pressure, i.e. increased ligand expression with unaltered receptor expression, at the maternal-fetal interface. Receptor-ligand interaction pairs are shown (Wilcoxon rank-sum test, p<0.05) using arrows from ligands towards receptors. Dot colours denote cell states/types, ligand squares illustrate average log_2_FC between conditions (only upregulated candidates are included from Logistic Regression; log_2_FC>0.25 and p-value<0.01 after Bonferroni correction) and receptors squares encode average expression. **(b,c)** vSTB-derived ligands *GDF15* and *INHBA* from (a) are validated in a multicentre cohort comparing gestational age corrected non-PE preterm controls with eoPE mRNA expression (n=23 eoPE, 11 preterm controls = pt.ctrl; sig.level>0.05; unpaired two-tailed t-test with Welch’s correction). **(d)** Immunofluorescence staining of GDF15 in l.ctrl and eoPE to validate differential mRNA-expression from (c) on protein level (n=3 each). **(e)** Whole slide scans with standardised exposure were evaluated using automated image analysis calculating mean GDF15 intensity on trophoblast area (n=3 each, scans divided into 4 quadrants; sig.-level>0.05, unpaired t-test). **(f)** *EP300* targets that are senescence-associated secretory genes differentially expressed in STB cell states in eoPE (heatmap showing log2-foldchanges with sig.-level>0.05; Logistic Regression and after Bonferroni adjustment for covariates) **(i)** Targeted high-resolution spatial transcriptomic data was acquired through spatially resolved *in situ* sequencing (ISS), with spatial context of mRNA molecules implicating highly structured tissue in an unsupervised 2D embedding (n=3 total, early and late villous samples; representative image of e.ctrl shown) **(j)** Senescence-associated INHBA expression is spatially variable in ISS *in situ* sequencing data and **(k)** is significantly more frequently found in proximity to vascular transcripts (black arrows). **(l)** Study design for early gestational maternal serum used for eoPE-risk-prediction. **(m)** Activin A, synthesised by inhibin βα dimers that are encoded by *INHBA*, predicts eoPE in a conditional logistic regression model and shows that the currently used FMF algorithm underestimates eoPE-risk in women with high first-trimester activin A (eoPE: n=27; healthy term controls: n=49, matched for BMI, gestational age, maternal age).

Our next aim was to identify the localisation of senescence-associated molecules within the trophoblast wall of villi. To do this, we compared spatially resolved transcriptomes from healthy term controls vs eoPE using *in situ* sequencing data (**Fig. 5i**). Transcripts were computationally assigned as villous wall structure and then qualified to have their proximity to vascular markers (*CDH5, IDO2, KDR, TEK, ZEB1*), derived from vessels close to the trophoblast layer, evaluated (see **Extended Data Fig. 8**). One finding from *in situ* sequencing data was a divergence in the spatial expression of the senescence marker *INHBA*. In eoPE, *INHBA* was found more frequently in proximity to vascular transcripts, while in term controls, it was found to be significantly distant from vessels (l.ctrl; **Fig. 5j**). The overall increase of SASPs in vSTB, in combination with spatial disorganisation of these senescent vSTB, suggests that there are functional dysregulations in regions of maternal to fetal crosstalk where oxygen and nutrient transport across the STB from maternal blood to fetal circulation occur. Further evidence for the importance of *INHBA* in the development of eoPE comes from our multicentre cohort, whose villous tissue also exhibited increases in *INHBA* expression (**Fig. 5k**). The gene products uncovered in this study, as described above (**Fig. 4c**), and their role in vSTB transition (**Fig. 2h-k**) and maternal-fetal crosstalk (**Fig. 5i-j**), might indicate a potential of these factors as an early predictor of eoPE.

Uteroplacental tissue analyses have been limited to cross-sectional studies at late timepoints due to the increased maternal and fetal risk for adverse outcomes following interventions during pregnancy. To evaluate the early predictive potential and pathomechanistic relevance of our identified markers (**Fig. 5a**), we turned to a prospective longitudinal cohort spanning from first trimester to delivery where clinical outcome of eoPE was verified^46^. Women were recruited in the first trimester of pregnancy. A serum sample was taken before carrying out a risk assessment to predict PE using the ‘Fetal Maternal Foundation’ (FMF) algorithm^16^. We excluded women with comorbidities such as chronic hypertension or diabetes mellitus and proceeded to clinically match women that later developed eoPE to control patients 1:2 (27 cases und 49 controls after technical exclusion of five samples; **Fig. 5l**). We used a conditional logistic regression model without prior risk stratification via the FMF algorithm for eoPE-prediction, and found that activin A also significantly contributed to predict eoPE already early in pregnancy (p=0.0123; **Extended Data Fig. 10**). Clinically, uterine artery pulsatility index measurements (UtA-PI) are performed to determine adverse pregnancy outcome. ROC analyses of UtA-PI in combination with Activin A and GDF-15 had a higher predictive potential compared to UtA-PI alone (**Fig. 5m**). Importantly, our model showed that women who exhibited high circulating levels of activin A and GDF15 in the first trimester had their risk of developing eoPE underestimated by the FMF algorithm.

Our multi-omics results from later in gestation were the origin to bridge evidence from this pilot study in early pregnancy to a longitudinal understanding of the pathomechanism. We used senescence markers based on our late gestation data (*INHBA*/Activin A^47^ and GDF15) in early pregnancy to assess and predict the risk of a later development of eoPE.

Overall, our data delineates that a preeclamptic placenta syndrome starts early in pregnancy during CTB to STB differentiation and ends with premature senescence transferring the placental transition defect into a maternal syndrome.

## Discussion

The pathomechanisms that lead to eoPE and distinguish at-risk mothers from their healthy counterparts have so far been unclear. Here, we reconstruct the course of pregnancy where longitudinal sampling is limited, and link evidence from preeclampsia in later gestation to early pregnancy where underlying pathomechanisms are still unclear and diagnosis is not yet possible. Thereby, we present evidence that the preeclampsia syndrome arises from an early pregnancy disruption of trophoblast differentiation. We trace the faulty trophoblast differentiation to the specific event of transition from progenitor-like vCTB to post-fusion vSTB that create the fetal barrier to maternal circulation. Our model suggests that vSTBs undergo a accelerated premature differentiation associated with a dysregulation of the transcriptional co-activator p300. This drives the premature development of a senescence-associated secretory phenotype of vSTB subtypes that is specific to eoPE and impedes maternal-to-fetal crosstalk. Accordingly, we provide evidence that levels of senescence marker activin A and GDF15, detectable in serum at early stages of gestation, are contributing to a prediction-model to estimate risk for the later development of eoPE. In summary, this model defines preeclamptic disease as a response to altered syncytiotrophoblast lineage drivers, which result in a disturbance of placental processes of senescence translated to maternal circulation. This is in line with previous hypotheses that senescence and the shedding of necroptotic STB bodies might play key roles in PE^48, 49^ and arise from disturbed trophoblast differentiation. We could capture these STB bodies in decidua and transcriptionally profile them (dDSTB). These dDSTBs are highly GDF15^pos^ STB particles derived from villous tissue and are putatively carried by maternal blood into the maternal vasculature in PE. They are known to regulate transcription in target maternal endothelial cells^50^.

Early uteroplacental tissue analyses derived from terminations of pregnancy have been limited by the difficulty of predicting the future clinical course of a pregnancy. To overcome this issue, we compared tissues obtained from a window at the beginning of pregnancy, at which time the further progression was unknown, with a late window, when the disease has fully developed. We had access to a longitudinal cohort that phenotyped women from their screening in the first trimester all the way to delivery. By comparing outcomes of women with eoPE to healthy controls, we could computationally extend findings of late pregnancy back to an early gestational window prior to the appearance of overt symptoms of the disease. Further computational work will need to be done that integrates biospecimens across a longer gestational time to interpolate further important time frames in PE development and progression.

We identified an early dysregulation in pre-eclampsia where p300-linked transcription factor complexes may affect the defective regulation of trophoblast development that is associated with STB fusion. Congruently, we identified upregulation of genes that are unique for a pre-fusion vCTB (CTBpf) and involved in fusion itself, such as *EVRFRD-1* and *GREM2*. This might be explained by Hippo-based mechanotransduction of high-velocity arterial shear stress that occurs secondary to the high-resistance uterine arteries^51^ phenotypical of eoPE. The disrupted trajectory of STB subtypes we described may act in concert with other influences – oxidative stress, altered perfusion, and other factors known to contribute to eoPE^2, 14^ – to create a senescence-associated dysfunction of the maternal-fetal barrier. This means that eoPE may be marked by a cell fate diversion that increasingly drives vCTBs toward the STB fate (**Fig. 1e**), via an autocrine feedback loop that is very likely mediated by activin.

Our study also lays the groundwork for a novel understanding of the early pathophysiology of the early-onset subtype of PE. We carry out a precise analysis of the fetal-to-maternal translation, from STB in the villi wall to maternal vasculature on the cellular level. In general, our work confirms well-established biomarkers of eoPE such as sFlt1 (*FLT1*), *PAPPA2*, and *PGF*, and identified additional early pregnancy markers that interact with maternal vasculature and are activin-pathway associated SASP factors, such as GDF15 (*GDF15)* and activin A (*INHBA*). We have shown these factors are pathophysiology-relevant and imminent to trophoblast transition. Trophoblast differentiation is disrupted and as sequelae, premature senescence of STB with increased *GDF15* and *INHBA* expression ensues. While activin A was reported previously, we can now additionally extrapolate it’s central role in the pathomechanism of preeclampsia early on. GDF15 was shown to be predictive of PE in term pregnancy^52^, and while it is putatively associated with cardiometabolic disease^53^, we found a new role connecting this with early pregnancy STB differentiation and early pathomechanisms.

Finally, we envision that the comprehensive multi-omics data we analysed will lead to the discovery of further pathomechanistic biomarkers and ultimately the development of new curative pharmacological approaches at early timepoints in gestation. These new approaches may finally prevent a syndrome that begins during a vulnerable period of the life of the child, and continues to affect them their entire lives. So far, the established symptomatic pre-eclampsia treatments, related to blood pressure management and preterm delivery, are primarily beneficial for the mother but have adverse effects on the fetus, thus posing a public health concern. We suggest that future molecular phenotyping and precision medicine in pregnancy should focus on the disturbed fetal trophoblast transition to prevent the pathogenesis of the fetus and subsequently of the mother.

## Methods

### Patient samples

Tissue sampling was done in a multicentre-design. Patients were recruited in Berlin (German), Graz and Vienna (Austria) and Oslo (Norway). The studies were approved by each regional committee and described individually and headlined by the analysing method.

### First trimester tissue used for snRNA-sequencing

Placental and matching decidual tissue were collected from electively terminated pregnancies with informed consent of healthy individuals (gestational age 5 – 11 weeks). Exclusion criteria were maternal age under 18, maternal BMI >25 and self-reported maternal pathologies. Ethical approval was obtained from the Medical University of Graz Ethics Committee (31-019 ex 18/19; 26-132 ex 13/14). Immediately after surgical extraction, tissue was stored at 4°C in culture medium DMEM/F12 1:1, 1 g/dL glucose, Gibco®, Life Technologies (TM), Thermo Fisher Scientific, Vienna, Austria) and processed in no more than 4 hours. Amnion was removed and decidua dissected. Villous and decidual tissue were separately rinsed twice in cold (4°C) 0.9% NaCl to remove blood afterwards snap frozen in liquid nitrogen and stored at −80°C until processing. Patient characteristics can be found in Extended Data Table 1.

### Healthy term tissue used for snRNA-sequencing

Healthy term samples were collected immediately after delivery at the inpatient clinic of the Department of Obstetrics and Gynaecology, University Hospital Graz, Austria. The study was approved by the local Ethics committee at the Medical University of Graz (31-019 ex 18/19; 26-132 ex 13/14) and informed consent was obtained from each participating woman. Representative tissue samples (1×1×1 cm) of the medial third of the placenta were cut from vital cotyledons that were macroscopically free of infarct areas or other obvious pathologies that are assumed to have happened during delivery. This should avoid sampling degraded RNA and ensure a high-quality yield for further analysis, well knowing that it might skew towards possibly inaccurate phenotypes on either side of disease and healthy samples. Amnion was dissected and remaining tissue were rinsed twice in cold (4°C) 0.9% NaCl to remove blood afterwards snap frozen in liquid nitrogen and stored at −80°C until processing. Patient characteristics can be found in Extended Data Table 1 and Supplementary Table 1.

### Early-onset pre-eclampsia and healthy term tissue used for snRNA-sequencing

Pregnant women were recruited in Oslo University Hospital prior to elective caesarean section after informed written consent, as previously described^1, 2^, from women with either early onset pre-eclampsia (eoPE) or normotensive pregnancies. eoPE was defined as new onset hypertension (blood pressure ≥140/90 mmHg) and new onset proteinuria (≥1+ on dipstick, or ≥30 protein/creatinine ratio) at ≥20 weeks gestation but with delivery prior to gestational week 34. Placental villous tissue biopsies were cut from the centre of central normal appearing cotyledons, and were snap frozen in liquid nitrogen and stored at −80°C until use. The study was approved by the regional committee for Medical and Health Research Ethics in South-Eastern Norway, and performed according to the Helsinki Declaration. Patient characteristics can be found in Extended Data Table 1 and Supplementary Table 1.

### Validation cohort Graz

Study samples were recruited retrospectively immediately after delivery at the inpatient clinic of the Department of Obstetrics and Gynaecology, University Hospital Graz, Austria between 2018 and 2019. Pre-eclampsia (PE) was defined according to the ISSHP guidelines (Brown MA, Pregnancy Hypertension, 2018). Women receiving low dose aspirin were excluded. The study was approved by the local Ethics committee at the Medical University of Graz (26-132 ex 13/14 and 31-019 ex 18/19) and informed consent was obtained from each participating woman. Patient characteristics can be found in Extended Data Table 2.

### Validation cohort Oslo

Pregnant women were recruited prior to elective caesarean section after informed written consent, as previously described^3^, from women with either pre-eclamptic (or normotensive pregnancies. PE was defined as new onset hypertension (blood pressure ≥140/90 mmHg) and new onset proteinuria (≥1+ on dipstick, or ≥30 protein/creatinine ratio) at ≥20 weeks gestation. In addition, eoPE was defined as delivery prior to gestational week 34. Placental villous tissue biopsies were cut from the centre of central normal appearing cotyledons, and were snap frozen in liquid nitrogen and stored at −80°C until use. The study was approved by the Regional committee for Medical and Health Research Ethics in South-Eastern Norway and performed according to the Helsinki Declaration. Patient characteristics can be found in Extended Data Table 2.

### Validation cohort Berlin

Samples from 19 placentas <34 weeks were collected from March 2013 to July 2014 at the Department of Obstetrics at Charité University Medicine, Campus Virchow Clinic, Berlin, Germany. The trial protocol was approved by the local ethics committee, and written and informed consent was obtained from all participants. Women were recruited at the time of clinical admission. PE was defined according to the International Society for the Study of Hypertension in Pregnancy (ISSHP) 2000, as new onset hypertension of >140/90 mmHg at two occasions six hours apart, in combination with proteinuria of >300 mg/24 h or >2+ dip stick. This subset was part of a bigger cohort that was described in detail previously^4, 5^. Patient characteristics can be found in Extended Data Table 2.

### Single-nucleus sequencing (snRNA-Seq)

#### Nuclei capture, library generation, sequencing

Approximately 100-200 mg frozen placental and corresponding, separately sampled, decidual tissue was processed according to an optimised nuclei isolation protocol by Krishnaswami et al.^20^ Briefly, frozen tissue was disrupted with a pre-cooled glass Dounce in homogenisation buffer (1X NIM2 [1X protease inhibitor, 1 µM DDT, 250 mM sucrose, 25 mM KCl, 5 mM MgCl2, 10 mM pH8.0 Tris], 0.4 U/µL RNAseIn, 0.2 U/µL Superasin, 0.1% v/v Triton X-100) and filtered through a flow-cytometry (BD Falcon) tube with a 35 µm cell sieve cap. Homogenate was incubated in the dark, on ice, for two minutes with DAPI (5 µg/µL) and centrifuged for eight minutes (1,000xg, 4°C). Pellet was resuspended with staining buffer, transferred to a FACS-tube (BD Falcon) with a 35 µm cell-sieve cap and analysed using the BD FACS ARIA III flow cytometer using the BD FACSDiva software (BD Bioscience). After FACS sorting with a cut-off at 90% viable single nuclei, nuclei from the landing buffer (1% BSA, 0.2 U/µL RNAseIn) were counted using a digital counting chamber (Elvira) to achieve the concentration of 400-500 nuclei/µl and were loaded onto 10x Genomics Chromium chips. 10x Genomics single-index v2 and v3 libraries were prepared according to manufacturer’s instructions (Chromium Single Cell 3’ Kits v2 User Guide – CG00052, Chromium Single Cell 3’ Kits v3.1 Dual Index User Guide – CG000315). Libraries were sequenced on an Illumina HiSeq-4000 (pair-ended) aiming for a minimum coverage of 50,000 raw reads per nucleus.

#### Data pre-processing and quality control

The demultiplexing, processing, identification of Unique Molecular Identifiers (UMI) and barcode filtering of raw 3’ snRNA-Seq data was performed using Cell Ranger software (versions 3.0.2, 6.0.1 & 6.1.2) from 10x Genomics. Specifically, the SP014 (10X V2 chemistry), SP082 and SP136 batches (10X V3 chemistry) were processed with versions 3.0.2, 6.0.1, and 6.1.2 respectively.

The transcripts were aligned against the pre-built human reference genome GRCh38 premRNA version 3.0.0, which was built from the GRCh38 precompiled reference (https://cf.10xgenomics.com/supp/cell-exp/refdata-cellranger-GRCh38-3.0.0.tar.gz), and modified for use with snRNA-Seq data by extracting “transcripts” features from the gene model GTF and instead annotating these as “exon”, as described in the protocol defined by 10x Genomics (https://support.10xgenomics.com/single-cell-gene-expression/software/release-notes/build#grch38_3.0.0). Subsequently, systematic biases and empty droplets were modelled and removed by filtering counts due to contaminated ambient RNA reads and random barcode swapping using the remove-background function implemented in CellBender v0.2.0^2^. The total number of droplets was kept at 15,000, and a combined ambient and swapping model was used. Filtered expression matrices were loaded into python v3.7.9 and further processed using scanpy v1.8.2.

The post-quantification quality control was computed with the calculate_qc_metrics function in scanpy. Nuclei having fewer than 200 expressed genes or for which the total mitochondrial transcript expression was higher than 5% were excluded. Only those genes expressed in more than three nuclei were included. Data quality was assessed by plotting the number of unique molecular identifiers (UMIs) and total number of genes per sample. After quality control filtering, the samples were log-normalised to 10,000 reads using scanpy.

#### Data harmonization, clustering, and cell annotations for placenta

For the data harmonization of placenta samples, firstly, 6000 highly variable genes were computed using scanpy’s highly_variable_genes function, using the dispersion-based method (flavor=’seurat_v3’) and otherwise default parameters. The donor identifier was used as the key batch to minimize selection of batch-specific genes. Subsequently, the samples were integrated using a Bayesian variational inference model scVI v0.14.5 (based on stochastic optimization and deep neural networks architecture). Using, scvi.model.SCVI and get_latent_representation functions in scVI, a shared latent space of 15 dimensions for all placental single nuclei was inferred. Precisely, 128 nodes per hidden layer, 2 hidden layers used for encoder and decoder for the variational inference, and 0.1 drop-out rate was used. Zero-inflated negative binomial distribution (ZINB) was used to model gene expression. Apart from using donor_id (each sample) as batch key, further categorical covariates (10X library chemistry used, procurement centre of samples, gestational time) and continuous covariates (total counts, total number of genes with at least one positive count, percentage of mitochondrial expression, XIST counts per nucleus) were used to minimize the influence of unwanted technical variation in the cell typing.

The K-nearest neighbour graph was computed on the scVI inferred latent space using pp.neighrbors function in scanpy with k = 15 and otherwise default parameters. To further reduce the high dimensional latent spaces to 2D, visualization was generated using Uniform Manifold Approximation and Projection (UMAP). Particularly, the umap-learn v0.5.2 implementation in python was used and the maximum number of iterations was set to 500 (for better convergence) and random state to 0 (for reproducibility).

Cell-typing (annotations) was initially performed on the control placenta samples (both early and late gestation) based on robust and specific expression of marker genes. At first, clusters were identified in an unsupervised manner using Leiden community algorithm implemented in scanpy (with an initial resolution limit of 2) and initially annotated using marker genes extracted from literature plus top signatures obtained from EmpiricalBayes-method by model.differential_expression function in scVI and Seurat’s FindAllMarkers Logistic Regression (LR) method in Seurat. Leiden clusters lacking robust/specific biological markers were merged into the closest cluster. Thereafter, a LR classifier model (optimized by the stochastic gradient descent algorithm) implemented in Celltypist v0.2.0^6^ was trained based on our control cluster labels and was used to predict the cell annotations in diseased (eoPE) samples. A confusion-matrix was used to evaluate the performance of the classifier (predicted labels) given the known ground-truth (from Leiden clusters annotation). Spurious Leiden clusters mapping to a specific sample and lacking appropriate markers were removed. Particularly, a fibroblast (n=669) and erythroblast (n=930) subpopulation were excluded because they mapped solely to two specific early samples and hence, do not contribute to comparative cell typing. Additionally, 547 misclassified nuclei firmly clustering with vCTBp but also expressing high STB/EVT markers were excluded (with further help from the pseudotime analysis where these nuclei could not be modelled in a specific differentiation path). For further internal validation of cluster phenotype, we computed module scores using known markers list using the tl.score_genes function in scanpy. Finally, all the clusters assigned to a phenotype (label) were evaluated using robust and specific marker genes (described in the Differential expression analysis section; also refer to Extended Data Figure 2).

#### Data harmonization, clustering, and cell annotations for decidua

Similar to placenta, the top 6000 highly variable genes were computed using scanpy’s highly_variable_genes function using donor_id as the batch key. Here, the cell typing was initially performed on the 10X V2 samples (because they were sequenced earlier) by annotating unsupervised Leiden clusters based on robust and specific markers expression. Using the get_latent_representation function in scVI, a shared latent space of 10 dimensions was inferred keeping the other parameters same as used for the placenta. Like placenta, markers were extracted from literature as well as top signatures obtained from Bayes-method scVI model.differential_expression function and Seurat’s FindAllMarkers LR method. Leiden clusters mapping uniquely to a donor were excluded for the purpose of comparative cell typing. Subsequently, the cell labels were transferred to the 10X V3 samples using scANVI^7^.

In parallel to scANVI, a LR classifier model from Celltypist was trained using the annotated cluster labels and was used to predict the cell annotations in 10X V3 samples. A confusion-matrix was used to evaluate the performance of CellTypist classifier and scANVI (predicted labels). Ultimately, each cluster was inspected using biological markers knowledge and final decisions were made.

A cluster (initially annotated as stromal given its proximity to the DSC1/2 and consisting of 2911 nuclei) was later excluded because of the expression of conflicting markers such as *NOTUM, HPGD* and *HLA-G* (denoting EVT lineage). It also expressed certain macrophage genes and was difficult to classify. The CellTypist LR classifier assigned them a very low confidence score (∼0) indicating the cluster was very likely contaminated. Another cluster (initially thought of as NKT cells; 1119 nuclei) were removed because of high macrophage gene expression.

The final UMAP embedding stratifying the cellular hierarchies for decidua and villi are shown in figure 1b.

#### Evaluation of clustering robustness

To ensure the effective annotation of cell types/states, amortized Latent Dirichlet Allocation (LDA) implemented within scVI was used to find topic profiles for both tissues. Conceptually, a distinct cell types/state should map to a unique topic. Subclusters share the topics of the mother cluster and in additionally usually harbor a unique topic. For example, dNK1 and dNK2 have shared as well as unique topics. This modelling approach can also be used to identify potential doublets when cells exhibit multiple conflicting topics (mainly due to opposing lineage markers), similar to marker-based approaches used in other single cell studies of placenta and decidua^8^.

LDA was performed at several stages (initially using the number of Leiden clusters equal to the number of topics and ultimately, to the number of final labels) to see if the learned topics were mainly dominant in cells close together in the UMAP space. When a topic is dominant in multiple clusters in the UMAP, it is an indicative of similarity between the clusters despite being distant in the embedding. This might happen if the local relationships were not preserved beyond a certain threshold. In this way, the problematic clusters were confirmed to not map to unique/known topics and hence, removed from all downstream analysis. Additionally, it was also used as a quality check for the UMAP embedding.

#### Syncytiotrophoblast sub-clusters

For the first time, we detected significant heterogeneity in the vSTB group. Notably, we found an interesting state termed as juvenile STB that is marked by the expression of paternally imprinted *DLK1* (regulator of cell growth and differentiation), *SPARC, TMSB10,* and *ACTB* which indicate an association with extracellular matrix remodelling and promotion of changes to cell shape. Interestingly, the juvenile nuclei robustly express the secretory phenotype (characterized by several PSGs and maternally imprinted *TFPI2*) and a classical STB like profile (expression of *CGA, CYP19A1, KISS1, ADAM12, SDC1* and others) for which it was classified under vSTB group. vCTBp was considered as the trophoblast progenitor as they are actively cycling given the expression of genes like *MKI67, TOP2A, STMN1* and *CENPK/CENPE*. Notably, they exhibit robust expression of *YAP1, TEAD1, TP63, CCNA2, ITGA6*- all known for their roles as progenitor. vCTBpf is primarily fusogenic and is characterized by very specific markers such as *GREM2, ERVFRD-1, ERVV-1/2, OTUB2,* and *DYSF*. Placental *F13A1+/FGF13+* resident macrophages (Hofbauer cells, vHBC) uniquely express hyaluronan receptor *LYVE1* in the immune cell subset, suggested to maintain arterial tone and have pro-angiogenic functions. We additionally identify antigen presenting *HLA-DRA+* placenta associated maternal monocytes/macrophages (vPAMM) which are villi-associated and are extra-embryonic or maternal in origin. A cycling population of vHBC was identified (vHBCp) characterized by traditional HBC genes as well as proliferative genes like *MKI67* and *TOP2A*. The villi myocytes (vMC) were identified given their *AGTR1* expression. Other cell types were mapped using well known based on their marker genes (Supplementary Table 8).

#### Evaluation of integration performance

To quantify integration performance for both decidua and placenta, we employed metrics suitable for atlas level integration as discussed by Luecken et al.^9^

Firstly, an adjusted rand index (ARI) implemented in scikit-learn was computed to ensure that the cluster labels are independent of the sample information (scaled from 0 to 1 where values close to zero indicate our labels were not influenced by batches). The score for decidua (0.062) and placenta (0.044) indicate our labels were not affected by batches (donor). Adjusted mutual information (AMI) was additionally used to verify the above observation.

Importantly, an absolute silhouette score (ASW) of batch labels per cell-type were computed to measure batch-mixing (scaled from 0 to 1 where 1 indicates ideal batch mixing and 0 represents strongly separated batches). Since the batches are expected to integrate within a cell identity, the batch ASW was computed per cell type/state. The mean scores for placenta (0.863) and decidua (0.802) cell type/states indicate well batch-mixing alongside bio-conservation.

#### Differential expression analysis

Cell-type marker analyses for both decidua and placenta were performed using multi-variate LR generalized linear model implemented in Seurat’s FindAllMarkers() and were further internally validated using the empirical Bayes method in scVI model.differential_expression function.

In case of LR, the number of UMI, number of genes, and percentage of mitochondrial transcripts per nuclei were used as continuous covariates. Additionally, ∼disease (if a nucleus is from a control or eoPE sample) and library (10X V2 or V3 chemistry) were used as categorical covariates to minimize the effects of eoPE and libraries. Only those genes having a log fold-change cut-off of 0.25 and expressed in at least 25% of cells within each cluster were considered as a significant cell marker. An adjusted p-value cut-off was kept at 0.05 (after Bonferroni correction for multiple testing).

#### Differential analysis of disease markers and gestational age correction

To determine the differentially expressed genes for disease (eoPE) against late controls, the LR framework (implemented Seurat’s FindMarkers function) was applied to respective cell types/states. The number of UMI, gene counts, percentage of mitochondrial transcripts and percentage of sex-specific transcripts per nuclei were used as continuous covariates.

Importantly, a cell type/state specific preterm-score was calculated using the preterm vs term in labour significant genes^10^ having FDR <0.05 and used as a continuous covariate in the LR model. This was explicitly performed to prevent strong preterm specific effects in the analysis since eoPE arises 6-8 weeks prior to healthy term. Since no differential features was separately reported for the vSTB preterm, the other trophoblasts genes (vCTB) were used for correction. Additionally, SLIT2 & ROBO1 genes were included in the module-score based correction for as they are associated with risk for spontaneous preterm birth^11^.

Additionally, cell type/state specific labour signatures were considered for correction by extracting the term in-labour vs no labour differential genes from respective groups^10^ Only those genes having a log2 fold-change cut-off of 0.25 and expressed in at least 10% of cells within each group were reported as significant given adjusted p-value < 0.05 (Bonferonni corrected). Both up- and down-regulated genes were computed.

No cell type/state exhibited significant composition shift in eoPE relative to term controls (except for vHBC)- hence, down sampling was only performed for the vHBC.

For cell types such as vCCT, dEVT, vCTBp, vCTBpf, dDSTB, dPC, dBcells no analysis was performed owing to extreme sparsity in eoPE group. None of our samples were confounded with a major co-occurring disease.

#### Reconstruction of differentiation trajectories, lineage relationships and computation of pseudotime genes

To infer the cluster and lineage relationships between the different trophoblast cell types/states- STREAM v1.1. (https://github.com/pinellolab/STREAM) and diffusion pseudotime^12^ were used. Specifically, the trajectory inference was restricted to the early controls of the trophoblast cell types including vCTBp (progenitor), vCTB, vCTBpf, vSTBjuv, vSTB1/2 and vCCT. In the late term controls, there is a striking discrepancy in the cell-type composition given a massive increase of vSTB sub-populations which signifies degradation rather than differentiation.

At first, the scVI harmonized control data as subsetted for the relevant cell types and learned the trajectory principal graph using STREAM 1.1. Using previously computed latent variables, cells were clustered in the reduced UMAP space for recovering the main and possibly finer structures of trophoblast differentiation. Thereafter, the principal graph was inferred on the manifold learnt from the dimension_reduction function using the first six components. K-means clustering was used for the initial graph seeding using seed_elastic_principal_graph(). The elastic principal graphs are structured data approximators, consisting of each cell as a vertex interconnected by edges. The inference of this graph relied on a greedy optimization procedure based on which a minimum spanning tree (MST) was constructed using the Kruskal’s algorithm. No branch pruning or shifting of nodes were performed to obtain the optimal principal graph (Figure 2A and Extended Data Figure 4a).

Ultimately, the transition and leaf markers were computed for all lineage paths (vSTB, vCTB and vCCT) by considering MKI67 positive vCTBp as a root node (start of the pseudotime) respectively. The transition genes are dynamical in nature and calculated by considering fold change in average gene expression of the first 20% and the last 80% of the cells for an individual branch based on the inferred pseudotime. For the genes exhibiting log2 fold change cut-off of 0.20, further Spearman’s rank correlation was calculated between pseudotime and gene expression of all the cells along the individual branches (as implemented in STREAM’s detect_transition_markers function). Ultimately, genes above a specified correlation threshold (=0.35) were reported as transition genes. For leaf gene detection, the z-scores of all leaf branches were calculated based on the average gene expressions. Particularly, Kruskal–Wallis H-test followed by a post-hoc pairwise Conover’s test (as implemented in STREAM’s detect_leaf_markers function) was used for multiple comparisons of mean rank sums test among all leaf branches. A Z-score cut-off of 1, and p-value cut-off of 0.01 were used to identify the candidate leaf genes. The expression of highly robust cell fate markers along the pseudotime provided a strong validation for our trajectories (Extended figure 4d).

To further evaluate the lineage relationships and global transcriptomic similarity between different cell types (for trophoblast), Diffusion map analysis was performed that orders cells based on their transcriptomic similarity in a Markovian space. This method considers each cell to be represented by a Gaussian wave function and diffusion distances are based on a robust connectivity measure between cells which is estimated over all possible paths of a certain length between the cells. The Eigen functions of the Markovian transition probability matrix (diffusion components; DC1 and DC2) were used for low-dimensional representation and visualization of trophoblast data (Extended Data Figure 4b).

#### Receptor-Ligand interaction databases

An extremely important factor deciding the results of the R-L interaction study is the underlying database used. Two popular databases, CellChatDB and FANTOM5, were used that allowed identification of well-established interactions such as MIF-ACKR3/CXCR7 and INHBA-ENG/END, which were unique to CellChatDB and FANTOM5 respectively.

#### Receptor-Ligand interaction differential analysis of eoPE vs term controls

The differentially expressed ligand-receptor interactions were inferred using Connectome v1.0.1 (specifically, the differential connectomics pipeline). For the maternal-fetal interface, the strategy was to use only secretory ligands for vSTB groups that can practically cross the maternal-fetal barrier and can be in contact with the maternal blood (decidua) where it can influence the vessels. Only significantly differentially upregulated ligands (p-value 0.01 after adjusting for multiple testing) in eoPE relative to term controls exhibiting a log2FC cut-off of 0.25 and detected in a minimum of 10% of diseased cells were considered. It was assumed that once a ligand is activated (upregulated), it would bind the receptor (irrespective of the latter being differential or not). Biologically, we can describe such instance as ligand pressure (where ligand is high, but receptor is either non-differential or low; Figure 4C). Multivariate Logistic Regression was used for differential calculation and the statistics are consistent with our former described DEG test (including covariate corrections).

For the within tissue interaction map (decidua and villi interaction) using immune and endothelial cell types, LR and Connectome were used. The log2FC cut-off of 0.25 and ligands/receptors detected in a minimum of 10% of diseased cells were considered for both up and downregulated differential candidates. The p-value was adjusted for covariates (as described for eoPE vs late term DEG). Visualization was performed using circos plots.

#### Receptor-Ligand interaction analysis in eoPE

The analysis for decidual STB and EVT ligands (with maternal VEC and dSMC) were restricted to eoPE samples only given their extreme sparsity in late term. Interactions were derived using Connectome using both FANTOM5 and CellChatDB databases. The min.cells.per.ident was kept at 75 and Diagnostic Odds Ratio (DOR) was calculated for each interaction pair. High DOR is an indicative of high specificity and sensitivity with a low rate of false positives and false negatives.

For dDSTB, the interaction list was filtered using pct.source (senders) >= 25% and pct.target (receivers) >= 20% (pct= percentage of cells expressing a ligand/receptor) and further, filtered by DOR.source > 3 and ligand expression > 1.5 (Figure 4A).

For the interactions with dEVT, relatively robust criteria were used for narrowing down the important interaction partners (from an initial list of > 2000 pairs). Particularly, DOR.source of 5, edge strength (product of the receptor and ligand expression) of 3 and minimum percentage of ligand expressing source of 50% was used to ensure cell specific communication.

#### Computational validation of major R-L interactions

All Connectome results were cross checked SingleCellSignalR^13^ for the vSTB, dDSTB and dEVT based interactions. All the R-L interactions were recapitulated (when not limited by database).

Subsequently, we applied additional tools (NATMI, logFC Mean (inspired by iTalk), CellphoneDB, CellChat) implemented within LIANA framework^14^ and we were able to recapitulate all R-L interactions across numerous databases. Identification of decidual and villous cell types and states.

#### Pathway and network analyses of marker genes

The list of DEG based on cut-off values (logFC +/-0.25 and a significance level of 0.05) were used as background for networks. Variable genes were excluded using the webtool diVenn (divenn.noble.org). Genes were then used as input in the stringDB for PPI networks (confidence level = 0.15, no added proteins in shells). Networks were then further analysed in Cytoscape (version 3.8.2)^15^. Hub genes, defined as genes with high correlation in candidate modules, were identified from the PPI network calculating top 5 genes of all topological analysis methods of CytoHubba^16^ in Cytoscape plug-in (DMNC; MNC, MCC, ecCentricity, Bottleneck, Degree, EPC, Closeness). The candidate hub genes were merged into one network, decomposed into communities using clustermaker^17^ and GLay^18^ Cytoscape plug-in based on Newman and Girvan’s edge-betweenness algorithm. The hub network was analysed to visualise the network degree of nodes by size of nodes. The original background logFC was used for continuous mapping colours. The hub gene network was used to calculate transcription factors via iRegulon^19^ cytoscape plug-in (standard threshold: enrichment score threshold 3.0, ROC threshold for AUC calc 0.03, Rank threshold 5000, minimum identity between orthologous genes: 0.0, max FDR on motif similarity: 0.001). Predicted transcription factors were visualised as PPI (confidence level 0.15) via stringDB and validated by adding expression values from the DEG list.

Pathway analyses are based on these DEG lists, hub genes, and transcriptions factors and were carried out via web-tools Metascape and Enrichr.

### *In Situ* Sequencing

#### High sensitivity library preparation

Fresh tissue samples of early villi were FFPE processed and stored at +4°C. A custom gene panel was used to detect specific cell-type and cell pathway genes of interest. The *in situ* sequencing method was processed according to manufacturer instructions (Cartana, part of 10x Genomics). 5µm tissue sections were baked at 60°C for one hour, deparaffinised in xylene, rehydrated in 100% and 70% ethanol, and permeabilised using citrate buffer (pH 6) for 45min at >95°C in a steamer. Sections were dehydrated in an ethanol series from 70 to 100% and air-dried (Secure Seal, Grace Biolabs, Bend, United States). Gene specific chimeric padlock probes were added, directly hybridised to the RNA at 37°C in an RNAse free humid chamber overnight and ligated at 37°C for 2 hours. Ligation derived circular oligonucleotide structures (padlocks) amplified overnight at 30°C. RNA-degradation during tissue processing was minimised by adding 0.1% v/v diethyl pyrocarbonate (DEPC) to all buffers and reagents not provided by the manufacturer.

#### Imaging

Imaging was performed using a digital slide scanner (Olympus SLIDEVIEW VS200) connected to external LED source (Excelitas Technologies, X-Cite Xylis). Fluorescence filters cubes and wheels were equipped with a pentafilter (AHF, excitations: 352-404 nm, 460-488 nm, 542-566 nm, 626-644 nm, 721-749 nm; emissions 416-452 nm, 500-530 nm, 579-611 nm, 665-705 nm, 767-849 nm) and single cube filters (Kromnigon; SpectraSplit 440, SpectraSplit 488, SpectraSplit Cy3, and SpectraSplit 594). Images were obtained with a sCMOS camera (2304 × 2304, ORCA-Fusion C14440-20UP, 16 bit, Hamamatsu), and Olympus Universal-Plansuperapochromat 40× (0.95 NA/air, UPLXAPO40X). To avoid signal cross-talk, the pentafilter was used to image DAPI, Cy5 and AF750 signals, and the single cubes to image AF488 and Cy3 were used. Imaged regions were recorded to perform repetitive cycle imaging. After imaging, labelling mix was stripped from each slide by adding three times 100% formamide for 1min, followed by a washing step.

#### Hybridizing and Sequencing

*In situ* sequencing steps were repeated six times with six different adapter probe pools, each imaged in five channels (DAPI, FITC, Cy3, Cy5, AF750). After stripping, adapter probes were hybridised at 37°C for 1 h in a RNAse free humid chamber, washed and sequencing probes hybridised at 37°C for 30 min in a RNAse free humid chamber. Sections were washed, dehydrated in an ethanol series, air-dried, and mounted with SlowFade Gold Antifade Mountant (Thermo Fisher Scientific). Library preparation protocols were optimised for placental tissue using high (*MALAT1*) and low (*RPLP0*) control probes before using the final probe panels. Background without any adapter probe pool was imaged in 6 channels for autofluorescence subtraction.

#### Image analysis and spot calling

Imaging data was analysed with the custom pipeline provided by CARTANA that handles image processing and gene calling. All code was written in MATLAB and additionally a CellProfiler pipeline (v.2.1.1)^20^ was used, that includes the ImageJ plugins MultiStackReg, StackReg and TurboReg as previously described^21^. In short, TIFF images from all sequencing cycles were aligned to the general stain of library preparation, and split into multiple smaller images. The median intensity of all RCP signals of each channel was calculated with an additional CellProfiler pipeline (v.4.0.7)^20^, this value was used to normalise RCP signal intensities of each channel to a pixel intensity of 10,000. The received multiplication factor value for each channel was integrated in the CellProfiler pipeline and the background of each channel subtracted from each sequencing cycle, to reduce the autofluorescence of the tissue. A pseudo-anchor was created for each cycle by making a composition of the four readout detection probe channels into one merged image. The pseudo-anchor was used to perform a second, more exact alignment. RCPs of the labelling mix were detected, x and y coordinates saved and fluorescence intensities measured. The highest intensity value in each sequencing cycle was assigned as positive event and used for decoding in Matlab. For signal visualisation, the selected transcripts were plotted on a DAPI-stained image.

### *In situ* sequencing data analysis

#### *In situ* sequencing data handling

*In situ* sequencing data handling was performed using the plankton.py v0.1.0 package (https://github.com/HiDiHlabs/planktonpy) in python 3.10.4. For the conclusive analysis routine, the in situ sequencing data was displayed as decoded spots of x and y coordinates of all detected mRNA molecules, each with an associated gene label. In total, three *in situ* sequencing slide scans were analysed (106KS, 107KS, 156KS). 156KS (early control) contained genes from the customized placenta/cell typing panel that was designed for retrieving cell and tissue types, and both 106KS (late control) and 107KS (eoPE) contained genes from the custom/pathway panel which was targeted at analysing cell state and metabolic activity (code book for panels available via zenodo doi: 10.5281/zenodo.5243240). The cell typing sample was taken during the early stage of pregnancy.

For visualization of the detected mRNA molecules in their histological context, matching DAPI stains of each sample slide were pre-processed by transforming it to grayscale, normalizing the colour values between 0 and 1 and pushing the low-exposure areas by raising all values to the power of 0.4.

#### Identification of cell type specific markers in the placenta panel

Analysis of the placenta cell typing data had the aim of contextualizing major cell types determined by snRNA-Seq analysis. Genes from the cell typing panel were conceptualized as cell type markers, with CTB, STB and HBC cell types considered for further spatial analysis and plotting since they had good marker coverage and constituted important spatial landmarks in the villi anatomy (with the walls being layered with STBs and CTBs, and HBCs forming distinct, compact cells in the intra-villous matrix).

Accordingly, a gene-cell-type affinity measure was derived through the gene molecule counts in the snRNA-Seq data set for CTB, STB and HBC. This was done per gene by contrasting molecule counts in the cells belonging to a cell type of interest in the snRNA-Seq data against the molecule counts of an opposing set of cells using plankton.py’s score_affinity function. Hence, for each analysis, two contrastive sets of cell types were defined: (i) CTB vs STB for CTBs; (ii) STBs vs CTBs for STBs; and (iii) HBC vs all other cells for HBCs. The transcriptome of CTB and STB cells could be expected to be more similar since both cells are trophoblasts. Therefore, to determine definitive cell type markers, CTB and STB were contrasted against each other, which would cancel out potential common trophoblast marker genes. Each genes’ mean molecule counts in all cells assigned to the two contrastive cell type sets was determined. The logarithm of the ratio of these mean count indicators was used as score for a gene’s affinity to a certain cell type. To improve visual clarity during plotting, a threshold of 0.5 was used to assign colour labels to each gene in both analyses. Genes exceeding this affinity score threshold determined markers for CTBs (*ASPM, ATAD2, BRIP1, CD24, CDH1, CENPE, DIAPH3, FBN2, KANK1, SEMA6D, TIMP3*), STBs (*ADAMTS20, CGA, CYP19A1, ENTPD1, KISS1, KLRD1, LEP, LINC00474, MYCNUT, PAPPA2, PLAC4, PLXDC2*), and HBCs (*CD163L1, CD36, F13A1, FGF13, LYVE1, MEF2C, SPP1*), with the remaining genes assigned as ‘other’.

#### mRNA molecule spatial context analysis in the placenta panel

Using the plankton.py’s run_umap function, a weighted neighbourhood graph was built using the 800 nearest neighbours of each molecule, with neighbours weighted according to their Euclidean distance using a Gaussian probability density function (PDF) at a bandwidth of 9 µm, which would roughly cover the area of a single cell and its immediate environment. Then, a model of local mRNA distribution was created over all genes by summing over each gene’s molecules’ weights. Furthermore, a regularization mechanism was introduced by increasing each distribution’s value of the gene of its molecule of origin by 1.15.

A 2D embedding of recurring spatial context was determined by applying the python umap-learn v0.5.3 algorithm to the local distributions. The number of neighbours in the UMAP algorithm was set to 24 and the minimal distance of points in the embedded representation was set to 0.2. UMAP used a Euclidean distance metric and was initiated at a random state of 42. The determined gene-celltype associations were used in a cell type visualization plot of the early placenta sample 156KS. All molecules were plotted as a scatter plot on top of the greyscale renderings of the DAPI stain. Molecules with a celltype-association affinity score above 0.5 were coloured accordingly, while the remaining molecules were rendered as grey.

#### Villi wall detection

Having demonstrated the principal plausibility of the spatial information in our in situ sequencing data during the cell-typing analysis, our experimental design required a follow up comparative analysis of pathways between a late control and an eoPE sample. The pathway categories chosen for the analysis of this second experiment were ‘vascularization’ (genes *IDO2, ZEB1, TEK, CDH5, KDR*), senescence (genes *MMP11, INHBA*) and trophoblasts (genes *LGR5, FGFR2, MET*).

Spatial analysis was restricted to the densely populated and well-structured villi walls in both samples, as this is the most structured part of the tissue. Villi walls were determined using a basic edge detection algorithm applied to matching DAPI signal, where villi walls were clearly remarked by dense nucleation. A greyscale rendering of the DAPI stain was smoothed using an optical Gaussian filter at a 2 µm bandwidth. Scikit-image’s (v0.19.2) feature.canny() implementation of the canny edge detection algorithm (using a sigma value of 3.7) was used to extract the villi walls in the smoothed image. Every molecule within a radius of 5 µm to any point of the detected edges was defined as being part of the villi walls, and all other molecules were discarded from further analysis. The wall filter algorithm was visualized by plotting the underlying DAPI stain in matplotlib’s violet-blue ‘magma’ colour scheme. The detected edges from the second step of the wall filter algorithm were plotted on top of the stain as orange lines. In the bottom-right half of the plot, the present mRNA molecules were plotted, coloured green or blue according to their assignment as wall/not-wall members (Extended Figure 6d).

#### Spatial relationship of vascular and senescence markers

For visualization of the spatial senescence-vascularization relation, all ‘wall’ molecules were plotted on top of a black-and-white rendering of the DAPI stain, coloured according to their gene assignments of ‘vascularization’ (red) and ‘senescence’ (yellow), with the other molecules plotted in a white. Optical inspection of the scattered senescence and vascularization markers hinted that senescence marker topography was more structured in the control sample as compared to the eoPE sample, with senescence marker expression being reduced around vascularization clusters in the tissue.

To statistically model this observation, the villi wall molecules were subdivided into two categories depending on their location of expression: (i) a vessel proximal category that contained all molecules within regions of 5 µm of another vessel marker; and (ii) a vessel-distal category that contained the other molecules. A null hypothesis was formed, according to which the distributions of genes should be equal within the two categories. For each gene, we reported the p-value of the violation of this null hypothesis in a binomial test. Scipy’s (v1.8.1) stats.binom.cdf implementation was used, with parameters ‘p’ defined as the total percentage of ‘proximal’ molecules, ‘k’ the gene-specific number of proximal molecules and ‘n’ the total number of molecules of the respective gene in the sample. The sorted p-values for all genes present in the pathway sample were displayed in a vertical bar graph, with the bars coloured according to their membership to the categories ‘senescence’, ‘vascularization’, ‘trophoblast’ or ‘other’ (Extended Data Figure 6f,g). The p-values of senescence, vascularization and a control category of ‘trophoblasts’ were extracted and plotted per sample as scatters on a vertical line. The scatters of both samples were displayed next to each other for visual comparison (Figure 4j, Extended Data Figure 6h).

### 10X Visium

#### Sample preparation

First trimester tissue was collected as described above, dissected under a stereomicroscope and snap-frozen by using isopentane in a liquid nitrogen bath. To avoid large batch-effects, multiple placental tissue sections were embedded in a 6.5×6.5 mm cryo-mould using OTC cryo-embedding medium (TissueTek). Samples were put on −80°C overnight and cryo-sectioned at −20°C. Control H&E staining was performed to ensure morphological intactness of the embedded tissues. The cryo-sectioned tissue at 10 µm was transferred to spatial transcriptomics slides (Visium, 10x Genomics) and placed on a single tissue optimisation and gene expression slide capture area.

#### Visium data sequencing

After having determined the permeabilisation time of 18 minutes following the tissue optimisation protocol (10x Genomics – CG000238 Rev A), the gene expression experiment was carried out according to the manufacturer’s user guide (10x Genomics – CG000239 Rev A). The images were scanned using the Slide Scanner Pannoramic MIDI (3DHISTECH) with the objective plan-apochromat 20×/0.8× (Zeiss).

The dual-indexed Visium library was then loaded at 200 pM and sequenced on a HiSeq-4000 (Illumina) with the following configuration: 28-10-10-90 (see sequencing requirements for Visium Spatial Gene Expression - https://support.10x_genomics.com/spatial-gene-expression/sequencing/doc/specifications-sequencing-requirements-for-visium-spatial-gene-expression).

#### Generation of the Visium count matrix

The base call (BCL) files generated from the Illumina run were converted to FASTQ reads with bcl2fastq (Illumina). Subsequently, the reads were mapped to the human reference dataset GRCh38 (build 2020-A; refdata-gex-GRCh38-2020-A) using the spaceranger count pipeline (Space Ranger v1.1.0) with automatic fiducial alignment and tissue detection. We observed 1,387 spots under the tissue, yielding 201,176 mean reads and 3,561 median genes per spot.

#### Seurat processing of the Visium count matrix

The 10x output folder was read using Load10X_Spatial function implemented in Seurat (v3). The object was then normalised with SCTransform function. PCA was then calculated using RunPCA: 50 PCs were computed and the first 20 were selected for identifying the k-nearest neighbours of each spot with FindNeighbors function. Finally, clustering was performed via FindClusters (resolution = 0.2).

#### Spotlight based deconvolution of Visium data

The spotlight object was generated using the spotlight_deconvolution function in SPOTlight (Version 0.1.7) by supplying the early villi subset (from the placental single-nuclei data) as reference. The marker table for the nuclei clusters was initially generated based on Logistic Regression method implemented in Seurat as discussed before, and subsequently filtered to yield the best topic profile representative of each cell type found in the dataset.

The non-negative matrix factorisation (NMF - *nsNMF*) regression as well as Non-negative Least Squares (NNLS) regression were used for deconvolution as implemented in SPOTlight. Cells contributing to less than 1% of the spot composition were removed, *min_cont = 0.01*. The deconvoluted spots were assessed by investigating the topic profiles of the cell type (Extended Data Figure 1f) and the nature of individual topics within a cell type.

### *In-vitro* validations

#### First trimester CTB cell culture

Villous cytotrophoblasts (vCTBs) were isolated according to recently published protocols^22^. Precisely, placental tissue (6 – 8^th^ week of gestation) was cut from the chorionic membranes, further minced into small pieces, and subjected to three consecutive digestion steps at 37°C in Hanks balanced salt solution (HBSS, Gibco) containing 0.25% trypsin (Gibco) and 1.25 mg/ml DNAse I (Sigma-Aldrich) for 10 min, 15 min, and 15 min, respectively. Per ml tissue, 5 ml digestion solution was used. Each digestion was stopped using 10% FBS ([v/v] PAA Laboratories). Subsequently the cells were filtered through a 100-µm pore size cell strainer (BD Biosciences), and cells from the second and third digestion were pooled. Next, the cells were loaded onto Percoll gradients (10 – 70 % [v/v]) and vCTBs were collected between 35 – 50% of Percoll layers, pelleted, and washed twice with HBSS. Eventually, red blood cells (if present) were removed by incubating vCTBs with erythrocyte lyses buffer (155 mM NH_4_Cl, 10 mM KHCO_3_, 0.2 mM EDTA, pH 7.3) at room temperature for 5 min. Afterwards, vCTBs were washed twice with HBSS. Subsequently, vCTBs were either immediately frozen in cell banker 2 (2 – 5 x 10^6^ per ml; Zenoaq) and stored in liquid nitrogen for flow cytometry analyses or seeded onto fibronectin-coated (20 µg/ml, Millipore) 48-well dishes at a concentration of 2.5 x 10^5^ cells per cm^2^ in DMEM/F12 (Gibco) containing 10 % FBS and 0.05 mg/ml gentamycin (Gibco). After 2 hours, non-attached cells were removed and fresh culture medium was added supplemented with vehicles (ctrl), or 5 µM A8301 (Tocris). For siRNA experiments, vCTBs were transfected with ON-TARGETplus non-targeting siRNAs (siCTRL; D-001810-10-0020, Thermo Scientific/Dharmacon), or co-transfected with ON-TARGETplus *YAP1* and *WWTR1* (siYAP1/TAZ; L-012200-00-0005, L-016083-00-0005, Thermo Scientific/Dharmacon) using Lipofectamine RNAiMAX (Invitrogen) according to the instructions of the manufacturer. After two and four days, the culture medium was changed containing the respective supplements and siRNAs. At indicated time points, cells were washed with ice-cold PBS and lysed using PeqGold Trifast (PeqLab). For immunofluorescence analyses cells were washed with ice-cold PBS and fixed with 4% paraformaldehyde for 15 min at room temperature.

#### RNA isolation and RT-qPCR

Cell pellets or pulverized tissue were lysed in QIAzol lysis reagent (Qiagen, Austin, Texas). RNA was isolated according to the manufacturer’s instructions (AllPrep DNA/RNA/Protein Mini, Qiagen, Austin, Texas). RNA quality was determined using an Agilent 2100 Bioanalyzer (Agilent Technologies, Santa Clara, CA, USA). Quality check was followed by reverse transcription of 1 μg total RNA per reaction using High-Capacity cDNA Reverse Transcription Kit (Applied Biosystems, Foster City, CA, USA), according to the manufacturer’s manual. For Graz cohort, qPCR was performed with Blue S’Green qPCR Kit (Biozym, CityVienna, Austria) using a Bio-Rad CFX96 cycler. For all other qPCRs, the QuantStudio 3 Real-Time PCR System (Applied Biosystems) with either TaqMan Fast Universal PCR Master Mix or Fast SYBR Green Master Mix (both Thermo Fisher Scientific) were used. Primer and probes (see below in Table S1) were designed using Real-time PCR (TaqMan) Primer and Probes Design Tool (online tool) from GenScript and synthesized by BioTez, Germany. Primers were diluted to a final concentration of 10 mM, probes to 5 mM. The target mRNA expression was quantitatively analysed with standard curve method. All expression values were normalized to the housekeeping gene *18S or TBP*. Validation cohorts were analysed individually and for combined presentation merged by z-transformation.

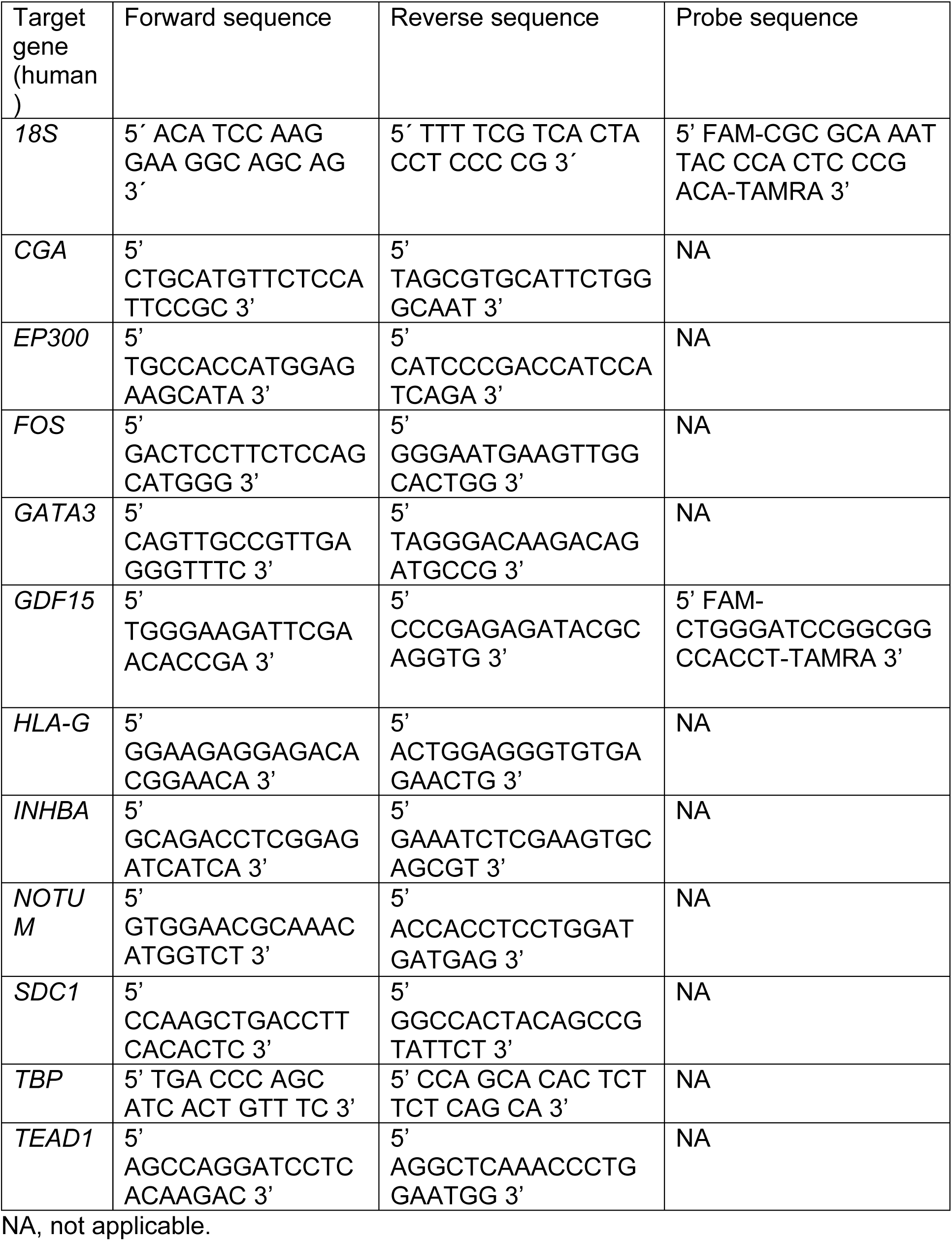

#### Flow cytometry analyses

Pooled villous cytotrophoblast cells from the same isolation (3 donors, gestational ages weeks 8+5, 9+2, 10+3) were seeded onto fibronectin coated plates (20 µg/mL, FC010 Merck) at a density of 0.25 x 10^6^ cells/cm^2^ and cultured for three (d3) or six (d6) days in DMEM/F12 (Gibco) containing 10% FBS and 0.05 mg/ml gentamycin (Gibco). Media was changed the day after thawing and every second day thereafter. d3 and d6 cells were trypsinised with TryPLE for 5 min at 37°C and plated on V plates alongside freshly thawed cells (d0) from the same donors at 0.1 x 10^6^ cells per well. All flow cytometry measurements included dead cell exclusion using Live Dead Fixable Aqua Dead Cell Stain Kit for 405 nm excitation (Thermo Fischer). ABTB2 (HPA020065) antibody was concentrated using the Antibody Concentration and Clean-Up Kit (Abcam, ab102778) and conjugated with PerCP-Cy5.5 (ab102911); GREM2 (ProteinTech, 13892-1-AP) was conjugated with Phycoerythrin (Pe) (ab102918), according to manufacturer’s instructions. Cells were stained with surface antibodies (Supplementary Table 2) in PBS + 0.5% BSA + 2 mM EDTA together with human FcR-blocking reagent (Miltenyi, 130-059-90) and incubated for 30 min on ice. Cells were stained with secondary antibodies (Invitrogen A11008 (anti-rabbit) & A-21235 (anti-mouse)) for 30 min on ice and fixed using the FoxP3 Staining Buffer Kit (eBioscience, 00-5523-00), stained with intracellular antibodies for 30 min on ice and analysed. Gating was performed according to unstained cells and fluorescence-minus-one (FMO) controls. Lineage-negative (CD34-, CD45-, CD49a-, CD235a-) cells were gated from live singlet events. Next, CD49f- (BD 747725) and E-cadherin+ (Cell Signalling 3195) cells were gated and co-expression of GREM2, CCR7 (BioLegend 353243) and ABTB2 in this subpopulation quantified. See extended Data Figure 6f for a visual representation of gates.

| **Antigen** | **Fluorophore** | **Clone** | **Dilution** | **Manufacturer** |
| --- | --- | --- | --- | --- |
| CD49f | BV421 | GoH3 | 1:200 | BD Biosciences |
| E-cadherin* | - | 24E10 | 1:50 | Cell signalling |
| GREM2<sup>^</sup> | PerPC-Cy5.5 | polyclonal | 1:200 | ProteinTech |
| ABTB2<sup>^</sup> | Pe | polyclonal | 1:100 | SigmaAldrich (HPA) |
| CCR7 | AlexaFluor 700 |  | 1:100 |  |
| CD34* | biotinylated | AC136 | 1:200 | Miltenyi |
| CD45* | biotinylated | REA747 | 1:100 | Miltenyi |
| CD49a* | biotinylated | REA1106 | 1:100 | Miltenyi |
| CD235a* | biotinylated | REA175 | 1:100 | Miltenyi |
| Streptavidin | PE-Vio 770 | - | 1:250 | Miltenyi |
| Secondary rabbit IgG | AF488 | - | 1:200 |  |

Antibodies used for flow cytometry. <sup>^</sup> intra-cellular antigen, \* non-preconjugated antibody, was stained with secondary streptavidin (if biotinylated) or anti-rabbit IgG.

#### First trimester Serum ELISA Measurement

Women were recruited in the first trimester of pregnancy and a serum sample was taken before risk assessment via the FMF algorithm^23^. We excluded women with comorbidities such as chronic hypertension or diabetes mellitus and proceeded to match women that developed early onset preeclampsia to controls 1:2 (n=28 vs n=56). The matching was based on the variables maternal age, gestational age at first scan, and BMI. This was done using the R package *Matching*, which finds for each case two matching controls that minimise the weighted distance of their matching variables. We excluded patients that were prescribed prophylactic Aspirin from being part of the control group to reduce confounding. The serum samples from the 84 selected case and control patients (matched on maternal age, GA at first scan, and BMI) were then analysed for leptin, perlecan, GDF15 and activin A according manufactures protocol: human HSPG (Perlecan) ELISA Kit (ab274393; Abcam), human GDF-15 Quantikine ELISA Kit (DGD150; R&D Systems), human/mouse/rat Activin A Quantikine ELISA Kit (DAC00; R&D Systems) and human Leptin Quantikine ELISA Kit (DLP00; R&D Systems). Due to missing samples and one sample that was removed after unreliable measurements, the group sizes were eoPE: n=27 and healthy term controls: n=49. A conditional logistic regression model was fit to the new data with predictor variables being included using forward selection. A conditional logistic regression model was calculated as absolute model without prior risk assessment based on the cohort published earlier^24^, a second model included the risk assessment by the FMF algorithm as offset. ROC curves and AUC are calculated from the method described in^25^, while confidence intervals stem from the DeLong method. R-scripts, data-tables and detailed results are available via https://github.com/HiDiHlabs/preeclamspsia_Nonn_etal/.

#### Immunofluorescence staining

Formalin fixed paraffin embedded (FFPE) placenta tissue sections (5 μm) were mounted on Superfrost Plus slides. Standard deparaffinisation was followed by antigen retrieval (AGR) in the multifunctional microwave tissue processor KOS in Tris-EDTA buffer pH 9.0 or citrate buffer pH 6.0 for 40 min at 93°C. Thereafter, sections were washed with PBS/T and incubated with Ultra V Block for 7 min at RT. For double staining, primary antibodies were mixed and diluted in antibody diluent and incubated on sections for 30 min at RT. Subsequently, slides were washed with PBS/T and incubated with secondary anti-mouse or anti-rabbit antibodies for 30 min at RT. Finally, slides were washed and nuclei stained with DAPI (1: 2,000; Invitrogen). Rabbit immunoglobulin fraction and negative control mouse IgG1 were used as described above and revealed no staining. Tissue sections were mounted with ProLong Gold antifade reagent (Invitrogen) and fluorescence micrographs were acquired with an Olympus microscope (BX3-CBH).

| Target | Clone | Species | Dilution | AGR | Company |
| --- | --- | --- | --- | --- | --- |
| Cytokeratin-7 | RCK105 (RUO) | mouse | 1:200 | pH 6.0 | BD Biosciences, 550507 |
| E-Cadherin | 4A2 | mouse | 1:500 | pH 9.0 | Cell Signaling, mAb #14472 |
| GREM2 | - | rabbit | 1:200 | pH 9.0 | proteintech, 13892-1-AP |
| p300 | D8Z4E | rabbit | 1:800 | pH 6.0 | Cell Signaling, mAb #86377 |
| anti-rabbit IgG (Alexa Fluor 555) |  | goat | 1:200 |  | Invitrogen |
| anti-mouse IgG (Alexa Fluor 488) |  | goat | 1:200 |  | Invitrogen |

#### Image analysis

Image analysis was performed on the whole-slide images using the image analysis software Visiopharm, version 2021.09. For p300 staining, we did villi and trophoblast detection and used the commercial Visiopharm app ‘Nuclei Detection, AI (Fluorescence)’ for nuclei detection and separation. We then classified nuclei into positive and negative by an intensity threshold of 70 (on an 8-bit scale 0-255) on the P300 marker within the respective nucleus area. Numbers of P300 positive and negative nuclei were assessed on the trophoblast area as well as on the remaining villous area.

For GDF15 staining, we accessed the mean intensity on the detected trophoblast area after villi and trophoblast area detection

#### Immunohistochemistry

Formalin fixed paraffin embedded (FFPE) placenta tissue sections were deparaffinised according to standard procedures. Antigen retrieval (AGR) was performed in a microwave oven in citrate buffer pH6 for 40 min. After a washing step with TBS/T sections were incubated with Hydrogen Peroxide Block (Epredia, Netherlands) to quench endogenous peroxidase followed by a further blocking step with UltraVision Protein Block (Epredia). Primary antibodies were diluted in antibody diluent and incubated on the sections for 45 min at RT. Slides were washed with TBS/T and thereafter the UltraVision LP HRP Polymer Detection System (Epredia) was used according to the manufacturer’s instructions. The polymer complex was visualized with AEC (AEC substrate kit, Abcam, UK), sections were counterstained with hemalaun and mounted with Kaisers glycerin gelatine (Merck, Germany). An Olympus VS200 slide scanner was used to scan the slides.

| Target | Clone | Species | Dilution | Company |
| --- | --- | --- | --- | --- |
| GDF15 | - | rabbit | 1:1000 (1<sup>st</sup> Trim)<br>1:250 (Term) | Sigma-Aldrich, #HPA011191 |
| HLA-G | 4H84 | mouse | 1:6000 | BD Biosciences, #557577 |
| βHCG | - | rabbit | 1:1000 | Thermo Scientific, #RB-059-A |

#### Spatial proteomics

Formalin fixed, paraffin embedded (FFPE) placenta tissue sections (5 μm) were mounted on PPS FrameSlides (Leica). Standard deparaffinisation was followed by antigen retrieval (AGR) in the incubator with Pepsin solution for 10 min at 37°C. Thereafter, sections were washed with PBS/T and incubated with Ultra V Block for 10 min at RT. For double staining, primary antibodies were mixed and diluted in antibody diluent and incubated on sections overnight at 4°C. Subsequently, slides were washed with PBS/T and incubated with secondary anti-mouse or anti-rabbit antibodies for 30 min at RT. Finally, slides were washed and tissue sections were mounted with Slow Fade Diamond Mounting media with DAPI (Invitrogen) and fluorescence micrographs were acquired.

| Target | Clone | Species | Dilution | AGR | Company |
| --- | --- | --- | --- | --- | --- |
| b-CG | 5H4-E2 | mouse | 1:300 | Pepsin | Abcam #ab9582 |
| E-Cadherin | 24E10 | rabbit | 1:200 | Pepsin | Cell Signaling, mAb #3195 |
| anti-rabbit IgG (Alexa Fluor 488) |  | goat | 1:300 |  | Life Technologies |
| anti-mouse IgG (Alexa in donkey Cy3) |  |  | 1:200 |  | Jackson Immuno Research |

Regions of interest were collected by laser microdissection (LMD) on a Leica LMD7 microscope using a 20x objective operated in fluorescence mode. An area of approximately 50,000 µm^2^ was collected per sample into 384-well plates (Eppendorf #0030129547). After LMD, tissue samples were processed for bottom-up LC-MS based proteomics as recently described^26^, but with small adjustments. Briefly, 4 µl of 60 mM triethylammonium bicarbonate (TEAB, Sigma #T7408) was added to each well, shortly centrifuged (2,000xg, 1 min) and the plate heated at 95°C for 60 min in a thermal cycler (Biorad’s S1000 with 384-well reaction module) at a constant lid temperature of 110°C. 1 µl of ACN was then added to each well (20% final concentration) and heated again at 75°C for 60 min in the thermal cycler. Samples were shortly cooled to room temperature and 2 µl LysC (Promega) added pre-diluted in ultra-pure water to 2 ng/µl and digested for 4 h at 37°C in the thermal cycler. Subsequently, 2 µl trypsin (Promega Trypsin Gold) was added pre-diluted in ultra-pure water to 2 ng/µl and incubated overnight at 37°C in the thermal cycler. Next day, digestion was stopped by adding trifluoroacetic acid (TFA, final concentration 1% v/v) and samples vacuum-dried (approx. 60min at 60°C). Finally, 4 µl MS loading buffer (3% acetonitrile in 0.2% TFA) was added, the plate vortexed for 10 s and centrifuged for 5 min at 2,000xg. Samples were stored at −20°C until LC-MS analysis.

#### LC-MS analysis

Liquid chromatography mass spectrometry (LC-MS) analysis was performed with an EASY-nLC-1200 system (Thermo Fisher Scientific) connected to a trapped ion mobility spectrometry quadrupole time-of-flight mass spectrometer (timsTOF SCP, Bruker Daltonik GmbH, Germany) with a nano-electrospray ion source (Captive spray, Bruker Daltonik GmbH). The autosampler was configured for sample pick-up from 384-well plates.

Peptides were loaded on a 20 cm in-house packed HPLC-column (75 µm inner diameter packed with 1.9 µm ReproSilPur C18-AQ silica beads, Dr. Maisch GmbH, Germany). Peptides separation followed a 32 min gradient with a flow rate of 250 nL with increasing concentration of buffer B (0.1% formic acid, 90% ACN in LC-MS grade H2O) to 60%. Buffer A consisted of 3% ACN, 0.1% formic acid in LC-MS grade H2O. The total gradient length was 44 min. Column temperature was controlled by a column oven and kept constant at 40°C.

Mass spectrometric analysis was performed in data-independent (diaPASEF) mode^27^ using the default method for long gradients with a cycle time of 1.8 s. Ion accumulation and ramp time in the dual TIMS analyser was set to 100 ms each and we analysed the ion mobility range from 1/K0 = 1.6 Vs cm-2 to 0.6 Vs cm-2. The total m/z range was set to 100-1,700 m/z. The collision energy was lowered linearly as a function of increasing mobility starting from 59 eV at 1/K0 = 1.6 VS cm-2 to 20 eV at 1/K0 = 0.6 Vs cm-2. Singly charged precursor ions were excluded with a polygon filter (timsControl software, Bruker Daltonik GmbH).

#### Data analysis of proteomic raw files

Proteomics measurements were analysed using the timsControl software (Bruker Daltonik GmbH, v. 3.1). For diaPASEF measurements, raw files were analysed with DIA-NN (v. 1.8)^28^ in library-free mode based on a predicted human spectral library (Uniprot 2021 release). Default settings were used with small adjustments. The mass range was set to 100 – 1,700 m/z, precursor charge state was 2 - 4 and the maximum number of allowed miscleavages was 2. MS1 and MS2 mass accuracies were set to 15 ppm and the match-between-runs option was enabled. Quantification strategy was set to ‘Robust LC’. For downstream data analysis, we used the protein FDR filtered pg.matrix.tsv and unique.genes.matrix.tsv DIA-NN output tables were analysed with Perseus (v. 1.6.15.0)^29^ and the Protigy R package (v. 1.0.2, https://github.com/broadinstitute/protigy). Missing values were imputed based on a normal distribution (width = 0.3; downshift = 1.8) after stringent data filtering (70% quantified values across samples). Prior to principal component analysis (PCA), batch effects were corrected with the proBatch R package (v. 1.10.0) based on the ComBat method (https://doi.org/10.3929/ethz-b-000307772). Pathway enrichment analysis was performed with clusterProfiler R package (version 4.2.2, https://doi.org/10.1089/omi.2011.0118).

#### Data and code availability

The snRNA-Seq raw data of the 33 villi and decidua samples generated in this study have been deposited in the European Genome-Phenome Archive under the accession number EGAS00001005681. The data are available under controlled access due to the sensitive nature of sequencing data, and access can be obtained by contacting the appropriate Data Access Committee listed for each dataset in the study. Access will be granted to commercial and non-commercial parties according to patient consent forms and data transfer agreements. Images of the ISS data are available via Zenodo (doi: 10.5281/zenodo.5243240). The Visium data are available via Zenodo (doi: 10.5281/zenodo.5336504). The remaining data are available within the article, Extended Data Figures or Tables and Supplementary Information. Scripts used to analyse the data and generate figures are available via https://github.com/HiDiHlabs/preeclamspsia_Nonn_etal/.

#### Data collection

No software was used for data collection.

#### Data analysis

Single-nucleus RNA sequencing analysis. The alignment and pre-processing of the snRNA-Seq data were performed using Cellranger version 3.0.2, 6.0.1 & 6.1.2. Ambient RNA and background noise correction were performed using CellBender 0.2.0. The data were processed using scanpy 1.8.2 in python 3.7.9. scvi-tools 0.14.5 was used for data harmonization. UMAP was computed using umap-learn 0.5.2. Trajectory analysis was performed using stream 1.1 and scanpy 1.8.2. Seurat 4.0 was used for marker analysis. Cell-cell interaction analyses were performed using Connectome 1.0.1 and LIANA 0.1.4. Gene/transcription factor regulatory network analyses and visualization were performed using STRING, iRegulon, and Cytoscape 3.8.2. For visualisation, igraph 1.3.2, circlize 0.4.15, dplyr 1.0.9, ComplexHeatmap 2.10.0; seaborn 0.10.0, and python-igraph 0.7.1 were used. Generally, scikit-learn 1.0.2, statsmodel 0.12.1, scipy 1.5.3, pandas 1.1.4, and numpy 1.19.4 were used.

#### 10X Visium was analyzed using Spotlight 1.0.0

ISS analysis was performed using python 3.10.4 and jupyter 1.0.0. Data handling was done using plankton 0.1.0, which uses pandas 1.4.3. All plots were generated using matplotlib 3.5.2. SnRNA-Seq data was integrated using scanpy 1.9.1. Image analysis for villi wall detection was performed with Scikit-image 0.19.2. Scikit-learn 1.1.1 was used to assign wall pixels and for spatial model building and nearest neighbor analysis. Numpy 1.22.4 was used for all algebraic operations on matrix representations of the data. Scipy 1.8.1 was used for statistical model building during pathway analysis.

Spatial proteomic analyses were performed using timsControl software (Bruker Daltonik GmbH, version 3.1), DIA-NN 1.8; and Protigy R 1.0.2, proBatch R 1.10.0, clusterProfiler 4.2.2, and Perseus 1.6.15.0. Pathway enrichment analysis was performed with clusterProfiler R 4.2.2 packages.

For conditional regression model analyses and visualisations, R 4.1.2, magrittr 2.0.2, Matching 4.9-11, tidyr 1.2.0, survival 3.2-13 and pROC 1.18.0 were used.

## Acknowledgements

We gratefully appreciate the excellent technical assistance of Juliane Anders, May-Britt Köhler, Gabriele N’Diaye, Jana Czychi, Kornelia Buttke (ECRC, MDC, Charité), and Sabine Maninger (Division of Cell Biology, Histology and Embryology, Medical University of Graz). We thank Russell Hodge (MDC) for proof reading the manuscript. We also thank Dr Andreas Glasner (Femina-Med) for the patient recruitment and technically outstanding sampling of first trimester placental material. We thank Daniel Kummer for the image analysis.

F. H. and D.N.M. were supported by Deutsche Forschungsgemeinschaft (HE6249/5-1; HE6249/7-1; HE6249/7-2; D.N.M.: Projektnummer 394046635 - SFB 1365) and BIH Omics platform, a joint focus area of the Berlin Institute of Health, Charité Universitätsmedizin Berlin and Max-Delbrück Center for Molecular Medicine. D.N.M. was supported by grants from the German Centre for Cardiovascular Research (DZHK; BER 1.1 VD). O. N. was supported through the PhD program Inflammatory Disorders in Pregnancy (DP-IDP) by the Austrian Science Fund (FWF): Doc 31-B26, PhD program Molecular Medicine at the Medical University of Graz, and through the Marietta Blau Grant by the Austrian Federal Ministry for Education, Science and Research (OeAD; BMBWF), and with grants from the MeFo Graz (PS-Stipendium 2019/2020), German Association of Prenatal Diagnostics and Obstetrics (DGPGM, 2020). K.S., T.K. and A.E-H. were supported by the K1 COMET Competence Center CBmed (Center for Biomarker Research in Medicine), which is funded by the Federal Ministry of Transport, Innovation and Technology (BMVIT), Land Steiermark (Department 12, Business and Innovation), BMWFW, the Styrian Business Promotion Agency (SFG), and the Vienna Business Agency. The COMET programme is executed by the Austrian Research Promotion Agency (FFG).

K.S. was supported by the Doctoral School in Translational Molecular and Cellular Biosciences at the Medical University of Graz.

S.H. and T.M. were supported by the Austrian Science Fund (FWF): P 34588

M.G. was supported by the FWF: (P 29639, P33554, I 3304, and Doc 31-B26) and the Medical University Graz through the PhD programs Inflammatory Disorders in Pregnancy (DP-IDP) and MolMed. B.H. was supported by the FWF: (Doc 31-B26) and the Medical University Graz through the PhD program Inflammatory Disorders in Pregnancy (DP-IDP). S.T. was supported by Federal Ministry of Education and Research of Germany in the framework of SAGE (031L0265). M.K. was supported by the Austrian Sciences Funds (P31470-B30). F.C and J.N. were supported by German Ministry of Education and Research (BMBF), as part of the National Research Node ‘Mass Spectrometry in Systems Medicine’ (MSCoreSys, grant agreement 161L0222). This work was supported by the BMBF-funded de.NBI Cloud within the German Network for Bioinformatics Infrastructure (de.NBI)

Schematic drawings were created with biorender.com.

## Author Information

The authors declare no competing financial interests. Correspondence and requests for materials should be addressed to.

## References

1. Williams, D. Long-term complications of preeclampsia. Semin Nephrol 31, 111–122, doi:10.1016/j.semnephrol.2010.10.010 (2011).

2. Brown, M. A. et al. Hypertensive Disorders of Pregnancy: ISSHP Classification, Diagnosis, and Management Recommendations for International Practice. Hypertension 72, 24–43, doi:10.1161/HYPERTENSIONAHA.117.10803 (2018).

3. Zeisler, H. et al. Predictive Value of the sFlt-1:PlGF Ratio in Women with Suspected Preeclampsia. N Engl J Med 374, 13–22, doi:10.1056/NEJMoa1414838 (2016).

4. Hauth, J. C. et al. Pregnancy outcomes in healthy nulliparas who developed hypertension. Calcium for Preeclampsia Prevention Study Group. Obstet Gynecol 95, 24–28, doi:10.1016/s0029-7844(99)00462-7 (2000).

5. Phipps, E. A., Thadhani, R., Benzing, T. & Karumanchi, S. A. Pre-eclampsia: pathogenesis, novel diagnostics and therapies. Nat Rev Nephrol 15, 275–289, doi:10.1038/s41581-019-0119-6 (2019).

6. Suryawanshi, H. et al. A single-cell survey of the human first-trimester placenta and decidua. Sci Adv 4, eaau4788, doi:10.1126/sciadv.aau4788 (2018).

7. Vento-Tormo, R. et al. Single-cell reconstruction of the early maternal-fetal interface in humans. Nature 563, 347–353, doi:10.1038/s41586-018-0698-6 (2018).

8. Liu, Y. et al. Single-cell RNA-seq reveals the diversity of trophoblast subtypes and patterns of differentiation in the human placenta. Cell Res 28, 819–832, doi:10.1038/s41422-018-0066-y (2018).

9. Garcia-Alonso, L. et al. Mapping the temporal and spatial dynamics of the human endometrium in vivo and in vitro. Nat Genet 53, 1698–1711, doi:10.1038/s41588-021-00972-2 (2021).

10. Birukov, A. et al. Blood Pressure and Angiogenic Markers in Pregnancy: Contributors to Pregnancy-Induced Hypertension and Offspring Cardiovascular Risk. Hypertension 76, 901–909, doi:10.1161/HYPERTENSIONAHA.119.13966 (2020).

11. Say, L. et al. Global causes of maternal death: a WHO systematic analysis. Lancet Glob Health 2, e323–333, doi:10.1016/S2214-109X(14)70227-X (2014).

12. Khan, K. S., Wojdyla, D., Say, L., Gulmezoglu, A. M. & Van Look, P. F. WHO analysis of causes of maternal death: a systematic review. Lancet 367, 1066–1074, doi:10.1016/S0140-6736(06)68397-9 (2006).

13. Lisonkova, S. & Joseph, K. S. Incidence of preeclampsia: risk factors and outcomes associated with early-versus late-onset disease. Am J Obstet Gynecol 209, 544 e541–544 e512, doi:10.1016/j.ajog.2013.08.019 (2013).

14. Rana, S., Lemoine, E., Granger, J. P. & Karumanchi, S. A. Preeclampsia: Pathophysiology, Challenges, and Perspectives. Circ Res 124, 1094–1112, doi:10.1161/CIRCRESAHA.118.313276 (2019).

15. O’Gorman, N. et al. Multicenter screening for pre-eclampsia by maternal factors and biomarkers at 11-13 weeks’ gestation: comparison with NICE guidelines and ACOG recommendations. Ultrasound Obstet Gynecol 49, 756–760, doi:10.1002/uog.17455 (2017).

16. Stern, C. et al. Low Dose Aspirin in high-risk pregnancies: The volatile effect of acetylsalicylic acid on the inhibition of platelets uncovered by G. Born’s light transmission aggregometry. J Reprod Immunol 145, 103320, doi:10.1016/j.jri.2021.103320 (2021).

17. Staff, A. C. The two-stage placental model of preeclampsia: An update. J Reprod Immunol 134–135, 1-10, doi:10.1016/j.jri.2019.07.004 (2019).

18. Roberts, J. M. & Escudero, C. The placenta in preeclampsia. Pregnancy Hypertens 2, 72–83, doi:10.1016/j.preghy.2012.01.001 (2012).

19. Gyllborg, D. et al. Hybridization-based in situ sequencing (HybISS) for spatially resolved transcriptomics in human and mouse brain tissue. Nucleic Acids Res 48, e112, doi:10.1093/nar/gkaa792 (2020).

20. Mund, A. et al. Deep Visual Proteomics defines single-cell identity and heterogeneity. Nat Biotechnol, doi:10.1038/s41587-022-01302-5 (2022).

21. Siddique, N. & Cox, B. Computational analysis identified accelerated senescence as a significant contribution to preeclampsia pathophysiology. Placenta 121, 70–78, doi:10.1016/j.placenta.2022.03.005 (2022).

22. Cox, L. S. & Redman, C. The role of cellular senescence in ageing of the placenta. Placenta 52, 139–145, doi:10.1016/j.placenta.2017.01.116 (2017).

23. Berns, D. S., DeNardo, L. A., Pederick, D. T. & Luo, L. Teneurin-3 controls topographic circuit assembly in the hippocampus. Nature 554, 328–333, doi:10.1038/nature25463 (2018).

24. MacDonald, T. M. et al. Circulating Delta-like homolog 1 (DLK1) at 36 weeks is correlated with birthweight and is of placental origin. Placenta 91, 24–30, doi:10.1016/j.placenta.2020.01.003 (2020).

25. Pique-Regi, R. et al. Single cell transcriptional signatures of the human placenta in term and preterm parturition. Elife 8, doi:10.7554/eLife.52004 (2019).

26. Knofler, M. & Pollheimer, J. Human placental trophoblast invasion and differentiation: a particular focus on Wnt signaling. Front Genet 4, 190, doi:10.3389/fgene.2013.00190 (2013).

27. Knofler, M. et al. Human placenta and trophoblast development: key molecular mechanisms and model systems. Cell Mol Life Sci 76, 3479–3496, doi:10.1007/s00018-019-03104-6 (2019).

28. Brosens, I., Pijnenborg, R., Vercruysse, L. & Romero, R. The “Great Obstetrical Syndromes” are associated with disorders of deep placentation. Am J Obstet Gynecol 204, 193–201, doi:10.1016/j.ajog.2010.08.009 (2011).

29. Zanconato, F. et al. Genome-wide association between YAP/TAZ/TEAD and AP-1 at enhancers drives oncogenic growth. Nat Cell Biol 17, 1218–1227, doi:10.1038/ncb3216 (2015).

30. Moya, I. M. & Halder, G. Hippo-YAP/TAZ signalling in organ regeneration and regenerative medicine. Nat Rev Mol Cell Biol 20, 211–226, doi:10.1038/s41580-018-0086-y (2019).

31. Okae, H. et al. Derivation of Human Trophoblast Stem Cells. Cell Stem Cell 22, 50–63 e56, doi:10.1016/j.stem.2017.11.004 (2018).

32. Haider, S. et al. Notch1 controls development of the extravillous trophoblast lineage in the human placenta. Proc Natl Acad Sci U S A 113, E7710–E7719, doi:10.1073/pnas.1612335113 (2016).

33. Gamage, T. K., Chamley, L. W. & James, J. L. Stem cell insights into human trophoblast lineage differentiation. Hum Reprod Update 23, 77–103, doi:10.1093/humupd/dmw026 (2016).

34. Rana, S., Burke, S. D. & Karumanchi, S. A. Imbalances in circulating angiogenic factors in the pathophysiology of preeclampsia and related disorders. Am J Obstet Gynecol 226, S1019–S1034, doi:10.1016/j.ajog.2020.10.022 (2022).

35. Roberts, J. M. et al. Preeclampsia: an endothelial cell disorder. Am J Obstet Gynecol 161, 1200–1204, doi:10.1016/0002-9378(89)90665-0 (1989).

36. Li, Y., Moretto-Zita, M., Leon-Garcia, S. & Parast, M. M. p63 inhibits extravillous trophoblast migration and maintains cells in a cytotrophoblast stem cell-like state. Am J Pathol 184, 3332–3343, doi:10.1016/j.ajpath.2014.08.006 (2014).

37. Meinhardt, G. et al. Pivotal role of the transcriptional co-activator YAP in trophoblast stemness of the developing human placenta. Proc Natl Acad Sci U S A 117, 13562–13570, doi:10.1073/pnas.2002630117 (2020).

38. Soncin, F. et al. Comparative analysis of mouse and human placentae across gestation reveals species-specific regulators of placental development. Development 145, doi:10.1242/dev.156273 (2018).

39. Salazar, V. S., Gamer, L. W. & Rosen, V. BMP signalling in skeletal development, disease and repair. Nat Rev Endocrinol 12, 203–221, doi:10.1038/nrendo.2016.12 (2016).

40. Herse, F. et al. Cytochrome P450 subfamily 2J polypeptide 2 expression and circulating epoxyeicosatrienoic metabolites in preeclampsia. Circulation 126, 2990–2999, doi:10.1161/CIRCULATIONAHA.112.127340 (2012).

41. Polesskaya, A. et al. CBP/p300 and muscle differentiation: no HAT, no muscle. EMBO J 20, 6816–6825, doi:10.1093/emboj/20.23.6816 (2001).

42. Wang, X. et al. P300 plays a role in p16(INK4a) expression and cell cycle arrest. Oncogene 27, 1894–1904, doi:10.1038/sj.onc.1210821 (2008).

43. Basisty, N. et al. A proteomic atlas of senescence-associated secretomes for aging biomarker development. PLoS Biol 18, e3000599, doi:10.1371/journal.pbio.3000599 (2020).

44. Bottner, M. et al. Characterization of the rat, mouse, and human genes of growth/differentiation factor-15/macrophage inhibiting cytokine-1 (GDF-15/MIC-1). Gene 237, 105–111, doi:10.1016/s0378-1119(99)00309-1 (1999).

45. Sugulle, M., Herse, F., Seiler, M., Dechend, R. & Staff, A. C. Cardiovascular risk markers in pregnancies complicated by diabetes mellitus or preeclampsia. Pregnancy Hypertens 2, 403–410, doi:10.1016/j.preghy.2012.03.002 (2012).

46. Guy, G. P. et al. Implementation of routine first trimester combined screening for pre-eclampsia: a clinical effectiveness study. BJOG 128, 149–156, doi:10.1111/1471-0528.16361 (2021).

47. Muttukrishna, S., Knight, P. G., Groome, N. P., Redman, C. W. & Ledger, W. L. Activin A and inhibin A as possible endocrine markers for pre-eclampsia. Lancet 349, 1285–1288, doi:10.1016/s0140-6736(96)09264-1 (1997).

48. Huppertz, B. Placental pathology in pregnancy complications. Thromb Res 127 Suppl 3, S96–99, doi:10.1016/S0049-3848(11)70026-3 (2011).

49. Hamad, R. R., Bremme, K., Kallner, A. & Sten-Linder, M. Increased levels of an apoptotic product in the sera from women with pre-eclampsia. Scand J Clin Lab Invest 69, 204–208, doi:10.1080/00365510802474384 (2009).

50. Wei, J. et al. Trophoblastic debris modifies endothelial cell transcriptome in vitro: a mechanism by which fetal cells might control maternal responses to pregnancy. Sci Rep 6, 30632, doi:10.1038/srep30632 (2016).

51. Leslie, K. et al. Increased apoptosis, altered oxygen signaling, and antioxidant defenses in first-trimester pregnancies with high-resistance uterine artery blood flow. Am J Pathol 185, 2731–2741, doi:10.1016/j.ajpath.2015.06.020 (2015).

52. Wong, G. P. et al. Circulating Activin A is elevated at 36 weeks’ gestation preceding a diagnosis of preeclampsia. Pregnancy Hypertens 27, 23–26, doi:10.1016/j.preghy.2021.11.006 (2022).

53. Wang, D. et al. GDF15: emerging biology and therapeutic applications for obesity and cardiometabolic disease. Nat Rev Endocrinol 17, 592–607, doi:10.1038/s41574-021-00529-7 (2021).

## References Methods

1. Harsem, N. K., Staff, A. C., He, L. & Roald, B. The decidual suction method: a new way of collecting decidual tissue for functional and morphological studies. Acta Obstet Gynecol Scand 83, 724–730, doi:10.1111/j.0001-6349.2004.00395.x (2004).

2. Staff, A. C., Ranheim, T., Khoury, J. & Henriksen, T. Increased contents of phospholipids, cholesterol, and lipid peroxides in decidua basalis in women with preeclampsia. American journal of obstetrics and gynecology 180, 587–592, doi:10.1016/s0002-9378(99)70259-0 (1999).

3. Herse, F. et al. Dysregulation of the circulating and tissue-based renin-angiotensin system in preeclampsia. Hypertension (Dallas, Tex. : 1979) 49, 604–611, doi:10.1161/01.HYP.0000257797.49289.71 (2007).

4. Hoeller, A. et al. Placental expression of sFlt-1 and PlGF in early preeclampsia vs. early IUGR vs. age-matched healthy pregnancies. Hypertens Pregnancy 36, 151–160, doi:10.1080/10641955.2016.1273363 (2017).

5. Ehrlich, L. et al. Increased placental sFlt-1 but unchanged PlGF expression in late-onset preeclampsia. Hypertens Pregnancy 36, 175–185, doi:10.1080/10641955.2017.1291673 (2017).

6. Dominguez Conde, C., et al. Cross-tissue immune cell analysis reveals tissue-specific features in humans. Science 376, eabl5197, doi:10.1126/science.abl5197 (2022).

7. Xu, C. et al. Probabilistic harmonization and annotation of single-cell transcriptomics data with deep generative models. Mol Syst Biol 17, e9620, doi:10.15252/msb.20209620 (2021).

8. Suryawanshi, H. et al. A single-cell survey of the human first-trimester placenta and decidua. Sci Adv 4, eaau4788, doi:10.1126/sciadv.aau4788 (2018).

9. Luecken, M. D. et al. Benchmarking atlas-level data integration in single-cell genomics. Nat Methods 19, 41–50, doi:10.1038/s41592-021-01336-8 (2022).

10. Pique-Regi, R. et al. Single cell transcriptional signatures of the human placenta in term and preterm parturition. Elife 8, doi:10.7554/eLife.52004 (2019).

11. Tiensuu, H. et al. Risk of spontaneous preterm birth and fetal growth associates with fetal SLIT2. PLoS Genet 15, e1008107, doi:10.1371/journal.pgen.1008107 (2019).

12. Haghverdi, L., Buttner, M., Wolf, F. A., Buettner, F. & Theis, F. J. Diffusion pseudotime robustly reconstructs lineage branching. Nat Methods 13, 845–848, doi:10.1038/nmeth.3971 (2016).

13. Cabello-Aguilar, S. et al. SingleCellSignalR: inference of intercellular networks from single-cell transcriptomics. Nucleic Acids Res 48, e55, doi:10.1093/nar/gkaa183 (2020).

14. Dimitrov, D. et al. Comparison of methods and resources for cell-cell communication inference from single-cell RNA-Seq data. Nat Commun 13, 3224, doi:10.1038/s41467-022-30755-0 (2022).

15. Shannon, P. et al. Cytoscape: a software environment for integrated models of biomolecular interaction networks. Genome Res 13, 2498–2504, doi:10.1101/gr.1239303 (2003).

16. Chin, C. H. et al. cytoHubba: identifying hub objects and sub-networks from complex interactome. BMC Syst Biol 8 Suppl 4, S11, doi:10.1186/1752-0509-8-S4-S11 (2014).

17. Morris, J. H. et al. clusterMaker: a multi-algorithm clustering plugin for Cytoscape. BMC Bioinformatics 12, 436, doi:10.1186/1471-2105-12-436 (2011).

18. Su, G., Kuchinsky, A., Morris, J. H., States, D. J. & Meng, F. GLay: community structure analysis of biological networks. Bioinformatics 26, 3135–3137, doi:10.1093/bioinformatics/btq596 (2010).

19. Janky, R. et al. iRegulon: from a gene list to a gene regulatory network using large motif and track collections. PLoS Comput Biol 10, e1003731, doi:10.1371/journal.pcbi.1003731 (2014).

20. Kamentsky, L. et al. Improved structure, function and compatibility for CellProfiler: modular high-throughput image analysis software. Bioinformatics 27, 1179–1180, doi:10.1093/bioinformatics/btr095 (2011).

21. Gyllborg, D. et al. Hybridization-based in situ sequencing (HybISS) for spatially resolved transcriptomics in human and mouse brain tissue. Nucleic Acids Res 48, e112, doi:10.1093/nar/gkaa792 (2020).

22. Haider, S. et al. Self-Renewing Trophoblast Organoids Recapitulate the Developmental Program of the Early Human Placenta. Stem Cell Reports 11, 537–551, doi:10.1016/j.stemcr.2018.07.004 (2018).

23. Guy, G. P. et al. Implementation of routine first trimester combined screening for pre-eclampsia: a clinical effectiveness study. BJOG 128, 149–156, doi:10.1111/1471-0528.16361 (2021).

24. O’Gorman, N. et al. Multicenter screening for pre-eclampsia by maternal factors and biomarkers at 11-13 weeks’ gestation: comparison with NICE guidelines and ACOG recommendations. Ultrasound in obstetrics & gynecology : the official journal of the International Society of Ultrasound in Obstetrics and Gynecology 49, 756–760, doi:10.1002/uog.17455 (2017).

25. Xu, H. et al. Estimating the receiver operating characteristic curve in matched case control studies. Stat Med 38, 437–451, doi:10.1002/sim.7986 (2019).

26. Mund, A. et al. Deep Visual Proteomics defines single-cell identity and heterogeneity. Nature biotechnology, doi:10.1038/s41587-022-01302-5 (2022).

27. Meier, F. et al. diaPASEF: parallel accumulation-serial fragmentation combined with data-independent acquisition. Nat Methods 17, 1229–1236, doi:10.1038/s41592-020-00998-0 (2020).

28. Demichev, V., Messner, C. B., Vernardis, S. I., Lilley, K. S. & Ralser, M. DIA-NN: neural networks and interference correction enable deep proteome coverage in high throughput. Nat Methods 17, 41–44, doi:10.1038/s41592-019-0638-x (2020).

29. Tyanova, S. et al. The Perseus computational platform for comprehensive analysis of (prote)omics data. Nat Methods 13, 731–740, doi:10.1038/nmeth.3901 (2016).

